# The largest radiations of freshwater fishes initiated at the Cretaceous-Paleogene boundary

**DOI:** 10.64898/2026.08.25.747131

**Authors:** Chase Brownstein, Bruno F Melo, Claudio F Oliveira, Thomas J Near

## Abstract

Freshwater biodiversity is disproportionally high relative to the limited availability of freshwater habitats. This pattern is exemplified by freshwater fishes. Over 50% of freshwater fish species are concentrated in a single clade, *Ostariophysi*, including the 5000 species of minnows, carps, and loaches, the 4500 species of catfishes, and the over 2000 species of tetras, pirahnas, and characins. However, the relationships and ages of ostariophysans remain uncertain. We show that the initial diversification of ostariophysans involved only two freshwater invasions and was driven by the strikingly rapid origination of major crown clades, including Neotropical electric fishes, lutefishes, and multiple major living clades of catfishes, carps and minnows, and tetras and characins, within five million years of the Cretaceous-Paleogene mass extinction. This result is congruent with the record of well-preserved body fossils of ostariophysans, but contrasts with the controversial assignment of isolated teeth and bones from the Cretaceous to nested lineages of this set of freshwater fish radiations. Although we confirm that *Alepocephaliformes*, an obscure marine lineage, is the living sister to *Ostariophysi*, our results demonstrate that the former clade only recently invaded the deep ocean, a transition that involved the loss of structures essential for enhanced auditory capabilities in ostariophysans and the related herrings and anchovies. These results establish a surprisingly young age for the major lineages of living freshwater fishes.

## 1. Introduction

Nearly 99% of water on Earth is contained in ice or the oceans, yet freshwater ecosystems harbor more species than marine environments [1,2]. This paradox of freshwater species richness is particularly striking in ray-finned fishes (*Actinopterygii*), which exhibit near-equal species diversity in freshwater and marine habitats [3–7]. However, freshwater ray-finned fish species are concentrated within a few lineages. Over half of all species in freshwater belong to the clade *Ostariophysi*, which includes catfishes, pacus, piranhas, tigerfishes and tetras, Neotropical electric fishes, and minnows, loaches, suckers, and carps [5,6,8–21]. The phylogenetic relationships among major ostariophysan lineages remain contentious despite the application of genomic sequence data to the problem [13,14,18–20,22–24], and the identity of their closest relatives are still unclear [11,12,25–27]. These uncertainties and the sparse early fossil record of ostariophysans [12,16,24,28–30] have obscured the timescale of ostariophysan diversification. It remains contentious whether ancient continental fragmentation events or marine dispersals and freshwater invasions are responsible for the origins of the more than 11,700 species in this clade [11,13,18,22,27,31–35].

Here, using a dataset of genomic sequences from 144 species representing all major lineages in the clade, as well as their potential sister lineages, the deep sea slickheads (*Alepocephaliformes*) and herrings, shads, and anchovies (*Clupeiformes*), we investigate the phylogenetic relationships and timescale of lineage origination in *Ostariophysi*. We show that *Alepocephaliformes* is the sister lineage to *Ostariophysi* and show that the deep sea and freshwater invasions that took place in these clades are associated with the loss and gain of hearing-related soft tissue and bony structures. Although we establish an Early Cretaceous origin and diversification of ostariophysans, our analyses suggest that continental endemism within major ostariophysan lineages results from rare Cenozoic dispersal episodes over the last 70 million years of Earth history. All major lineages of ostariophysans have crown ages that approximate or immediately postdate the Cretaceous-Paleogene mass extinction, implying that, contrary to expectations from the fossil record of freshwater faunas as a whole [36–40], these ecosystems experienced significant faunal changes after the asteroid impact 66.02 million years ago. Like mammals and birds, living lineages of ostariophysan fishes diversified in response to the ecological opportunities that emerged after the extinction of 75% of species at the end of the Mesozoic, and following their invasion of the freshwater realm.

## 2. Methods

*(a) Systematics.—*Here, we deploy the phylogenetic rank-free taxonomy of ray-finned fishes formalized by Near and Thacker [12] in accordance with the rules of the PhyloCode [41]. Following these conventions and in accordance with emerging trends in the literature [42], we italicize all clade names in this paper.

*(b) Sequence dataset assembly.—*To capture the phylogenetic diversity of ostariophysans and relatives, we sequenced ultraconserved elements (UCEs) from all major lineages of *Ostariophysi* and the larger clade *Otocephala* and combined these sequences with those harvested from published whole genomes to produce a dataset totaling 157 individuals representing 144 species with six outgroups (Table S1). Our analyses include the first genomic sequence data for several deeply divergent otocephalan lineages, including tubeshoulders (*Platytroctidae*), one of the two major lineages of *Alepocephaliformes* [43], and the gonorhynchiform genera *Kneria, Cromeria*, and *Grasseichthys*. We include new UCE sequence data from 77 species. Using previously published methods [13,14,31,32,44–50], we extracted DNA from tissues using DNeasy Blood and Tissue Kits (Qiagen), quantified DNA extractions using a Qubit fluorometer (Life Science Technologies), confirmed high-weight DNA using gel electrophoresis, and sent dried-down DNA plates for sequencing at Arbor Biosciences (Ann Arbor, MI) using the UCE probe set for *Ostariophysi* [14]. We used *phyluce* v. 1.7.3 [51] to search genomes for UCEs and combine them with those sequenced and produce sequence alignments using built-in commands from MAFFT v. 7.505 [52]. Next, we used *CIAlign* [53] to identify and remove chimeric UCE alignments. This approach retained averages of 1329 loci and 1,344,107 bp per individual. Our genome skimming approach only recovered a small number of loci for several non-ostariophysan sequences on GenBank. As a result, our 75% complete dataset sampling all 163 tips comprised 254 UCE loci. We tested whether the resolution of ostariophysan relationships depended on the size of our sequence dataset by subsampling 37 species for which over 1,100 UCEs were recovered that represent all major lineages in *Otocephala.* We then used these two datasets, hereafter called the ‘complete’ and ‘reduced’ datasets, for downstream analyses.

*(c) Phylogenetic Analyses and Concordance Factor Estimation.* We conducted all phylogenetic analyses under maximum likelihood in *IQ-TREE* 2 [54,55]. For both the complete and reduced datasets, we inferred maximum likelihood phylogenies using the concatenated sequence data under single and multiple partition schemes using PartitionFinder 2 [56,57] to select optimal partitioning schemes. We also inferred gene trees to use for species tree inference under the multispecies coalescent model implemented in ASTRAL-III v. 5.7.8 [58]. For all phylogenetic analyses, we used ModelFinder [59] to select best-fit models of nucleotide evolution, and assessed nodal support by using ultrafast bootstrapping over 1000 replicates. Finally, we used gene and site concordance factors [60], which measure the number of decisive gene trees and sequence sites, respectively, that support a given node in a species tree, to more thoroughly interrogate support for phylogenetic relationships.

*(d) Time Calibration.—*We time-calibrated our phylogeny of *Otocephala* using a Bayesian node-dating approach implemented in BEAST v. 2.6.7 [61,62] on three sets of 30 randomly sampled UCE alignments from the ‘complete’ set. Owing to uncertainties surrounding the phylogenetic positions of extinct species of ostariophysans, we compiled a set of fossil calibrations that we justified based on either the results of explicit phylogenetic analyses of morphological characters, or, in cases where phylogenetic analyses were not performed, the presence of unambiguous synapomorphies of particular clades. Our calibration list included 15 fossils (Supplementary Text). For all time-calibration analyses, we used a general time reversible model of nucleotide evolution with the +G gamma among-site rate variation parameter, a relaxed log-normal molecular clock, and the implementation of the Fossilized Birth-Death (FBD) Model in BEAST2 [63] as the branching model. The FBD model has the advantage of being identifiable [64] and allows for the inclusion of an extant sampling fraction and sampling parameter. To take advantage of this and also account for the uncertain placements of fossil stem-group species relative to one another, we used a node-dating protocol to time-calibrate our phylogeny under this branching model; this protocol has been shown to give overlapping age distributions to traditional use of the FBD model where multiple successive stem-lineage fossils are included as tips [65]. We specified the rho parameter of the FBD model as 0.01, which is the total number of ingroup species (n=144) in our dataset divided by the total number of valid otocephalan species (12683) listed in Eschmeyer’s Catalog of Fishes [4,66] in January of 2025. We set the diversification rate prior to 0.05, the approximate background diversification rate estimated for teleost fishes in previous analyses [13,50], with bounds of 0.00 and 1.00. We placed fossil calibrations at nodes using monophyletic lognormal MRCA priors that we adjusted so that 97.5% of the distribution of the age fell before the fossil occurrence in each case. The tree topology was fixed to the single-partition concatenated phylogeny inferred using the complete dataset. For each set of UCEs, we ran three independent MCMC chains over 200 million generations with a 100 million generation pre-burnin. We checked for convergence of the posteriors and effective sample sizes values over 200 using Tracer v. 1.7 [67]. Finally, we combined the top 25% of posterior tree sets in *LogCombiner* v. 2.6.7 subsampling every 5000 generations and annotated the posterior tree set to the target tree topology with median node heights in *TreeAnnotator* v. 2.6.6.

*(e) Historical Biogeographic Reconstructions.—*We conducted historical biogeographic reconstructions on the time-calibrated phylogeny of *Otocephala* using the R package BioGeoBEARS [68] using two area schemes following a recent study on characiphysans [69]. In one scheme, we coded areas by continents and oceans, for a total of five continents and three oceans. In the other, we coded species by their presence in the freshwaters of northern hemisphere (Laurasia), freshwaters of southern hemisphere (Gondwana), or in oceans and seas (Oceanic). Next, we fit three different biogeographic models with and without the addition of a jump dispersal parameter (+j). These were a simple dispersal-extinction-cladogenesis (DEC), a dispersal-vicariance-like (DIVALIKE) model, and Bayesian model (BAYAREALIKE). For each scheme, we compared the fit of these models using log-likelihood and weighted Akaike Information Criterion (AIC) scores and then selected the best fit model for historical biogeographic reconstruction. Finally, we compared the ages of inferred intercontinental dispersal and vicariant events to reconstructed land boundaries in two recently released sets of paleogeographic reconstructions [70,71].

*(f) Ancestral State Reconstructions of Habitat.—*We conducted all ancestral state reconstructions using the R package *phytools* v 2.0 [72,73]. We ran ancestral state constructions on two different coding schemes: one where habitat was treated as a binary trait (exclusively freshwater/brackish vs marine component) and one where habitat was considered a three-state, polymorphic character. For each, we coded species based on their corresponding species account in the FishBase database. Ancestral state reconstructions of aquatic habitat have varied in their treatment of this trait [24,74–80]. Because many species access multiple aquatic habitats, it is not sensible to code habitat as a discrete trait with strictly non-overlapping states [77]. To deal with this problem in the case where habitat was coded as a three-state trait, we treated habitat as a polymorphic character using the fitpolyMk function with unordered transition rates and a π root prior distribution [81]. For the analysis of habitat as a binary trait, we compared the fit of two models using AIC scores: an equal transition rates (“ERD”) model and an unordered rates (“ARD”) model. Following model fitting in each case, we conducted stochastic mapping over 1000 simulated topologies that we summarized in single ancestral state reconstructions.

*(g) Ancestral State Reconstructions of Auditory Structures.—*We searched the literature for information on the distribution of features relevant to increased auditory capacity in otocephalans, including the presence of a swimbladder, the Weberian apparatus, and the clupeiform gas-filled bullae. Next, we used *phytools* v. 2.0 [72,73] to test the fit of four models of character evolution: the ERD and ARD models, as well as irreversible models that forbade forward or backward transitions among character states in accordance with Dollo’s Law [82]. Finally, we ran stochastic mapping over 1000 simulated topologies for each trait and summarized the results in single ancestral state reconstructions.

## 3. Results

Our phylogenomic analyses maximizing both taxon and sequence sampling (Figure 1; Figure S1-S7) support the placement of *Clupeiformes* as the sister lineage to all other otocephalans, the placement of *Alepocephaliformes* as the living sister lineage to *Ostariophysi*, and the placement of lutefishes (*Cithariniformes*) outside of a clade containing tetras and piranhas (*Characiformes*) and catfishes (*Siluriformes*; Figure 1, Figures S1–S7)[12]. The relationships of *Alepocephaliformes* among ray-finned fishes are considered unstable [11,12,26,27] and have rarely been tested using more than a handful of loci [25,83]. Similarly, whether *Cithariniformes* and *Characiformes* form a clade has been the subject of much controversy [14,18–20,22,31]. Our phylogenomic analyses also demonstrate the reciprocal monophyly of both tubeshoulders (*Platytroctidae*) and slickheads (*Alepocephalidae*) in *Alepocephaliformes* for the first time using genome-wide marker data. We infer several relationships recently proposed using genomic sequence data with strong nodal support, including the placement of Denticle Herring *Denticeps clupeoides* as the living sister to other *Clupeiformes* [25,84], the placement of algae eaters (*Gyrinocheilus*) as the living sister to all other *Cypriniformes* [85,86], the monophyly of the recently recognized characiform clade *Lepidarchidae* and its placement within a larger Africa tetra and tigerfish radiation [32], the placement of the enigmatic leaf-litter dwelling fish *Tarumania walkerae* as sister to *Erythrinidae*, and the sister relationship of the South American knifefish lineages *Apteronotidae* and *Sternopygidae* (Figure 1; Figures S1-S7)[87] in the recently named clade *Sinusoidei* [88]. In *Siluriformes*, we inferred that armored catfishes (*Loricarioidei*) are the living sister to all other catfish lineages and *Pimelodoidea* as a clade nested deeply within the globally distributed *Siluroidei* (Figure 1; Figures S1-S7).

**Figure 1.**
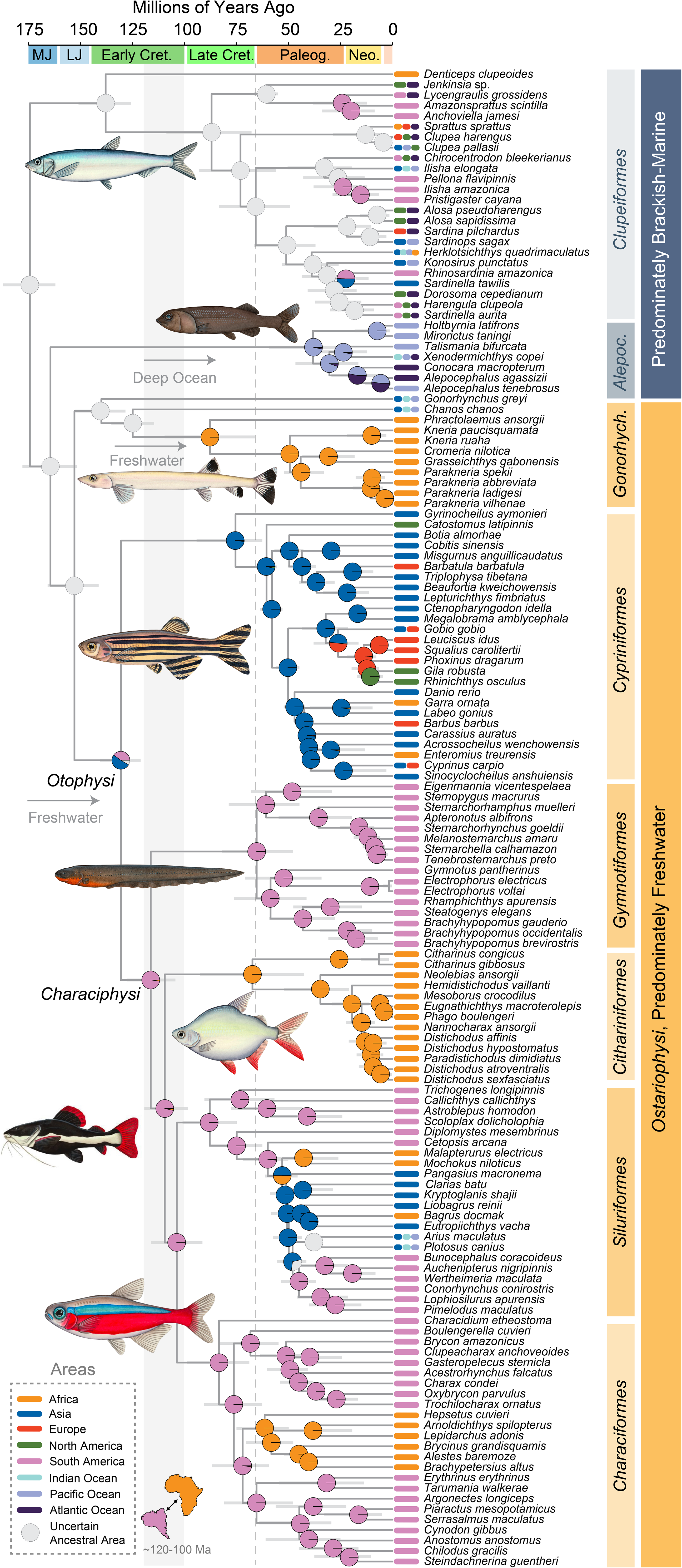
Evolutionary History of Ostariophysan Fishes and Their Relatives. Time-calibrated phylogeny from the Bayesian node-dating analysis of three sets of 30 UCE alignments of 144 otocephalan species conducted in BEAST v. 2.6.7. Bars at nodes indicate 95% highest posterior density (HPD) intervals for divergence times, and pie charts at nodes indicate probabilities for the node occupying particular areas in the historical biogeographic reconstruction conducted in BioGeoBEARS on the eight area dataset using the preferred model (BAYAREALIKE). Bars at tips indicate the present distributions of species among the areas in the eight area scheme. Gray-shaded region indicates the period when Africa and South America separated, and the dashed red line indicates the K-Pg boundary. Abbreviations: MJ, Middle Jurassic; LJ, Late Jurassic; Cret. Cretaceous; Paleog., Paleogene; Neo., Neogene; *Alepoc*., *Alepocephaliformes*; *Gonorhynch.*, *Gonorhynchiformes*. Illustrations are by Julia Johnson (lifesciencestudios.com).

Our relaxed molecular clock analysis (Figure 10 estimates a Late Jurassic origin for *Ostariophysi* [median most recent common ancestor (MRCA) age: 152.88 Ma, 95% highest posterior density intervals (HPD): 141.24, 165.78 Ma] following the origin of *Otocephala* in the latest Early Jurassic 174.32 Ma (95% HPD: 162.04, 187.18 Ma) and the most recent common ancestor of *Alepocephaliformes* and *Ostariophysi* at 164.43 Ma (95% HPD: 151.72, 177.88 Ma)(Figure 1, Figure 3). Despite the ancient divergence with *Ostariophysi*, the common ancestor of *Alepocephaliformes* is young, dating to the Late Eocene 38.22 Ma (95% HPD: 21.51, 59.37)(Figure 3).

All other divergences among major lineages of ostariophysans occur in the Early Cretaceous, between 135 and 100 million years ago (Figure 1). Notably, our historical biogeographic reconstructions do not unambiguously support any associations between the initial divergences in ostariophysans and the fragmentation of the supercontinents Laurasia and Gondwana (Figure 2). The divergence of the predominately northern hemisphere *Cypriniformes* from other lineages of *Otophysi*, all of which predominately occupy the southern hemisphere, occurs at 130.57 Ma (95% HPD: 121.06, 140.58 Ma), far postdating the separation of Laurasia and Gondwana by the Late Jurassic [70,71], though coinciding with the existence of a proposed Early Cretaceous dispersal corridor between the supercontinents through Europe [89–93]. A second vicariant event driven by continental fragmentation may explain the divergence of the exclusively African *Cithariniformes* from a clade containing *Siluriformes* and *Characiformes*, which occurs at 109.47 Ma (95% HPD: 98.25, 121.18 Ma), following the complete isolation of Africa from South America, which is unambiguously reconstructed as the area occupied by the common ancestor of these three lineages (Figure 1). However, the origination of crown *Cithariniformes* 30 million years later implies that a dispersal to Africa from South America could have occurred anytime during the Late Cretaceous, following the complete fragmentation of Gondwana.

**Figure 2.**
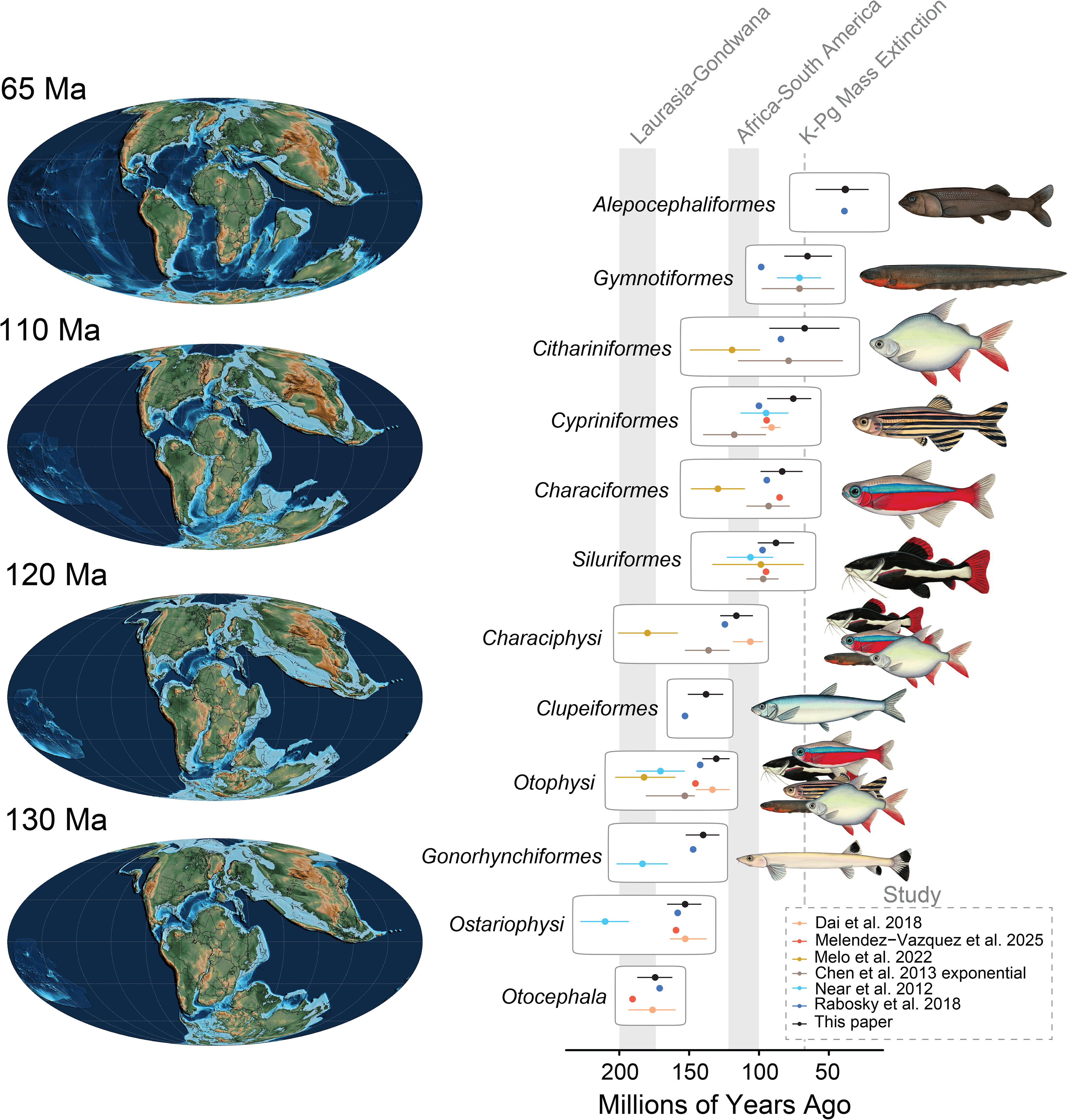
Ostariophysan Origins, Continental Fragmentation, and the K-Pg Mass Extinction. Mollweide paleogeographic maps from Scotese et al.[70] at left depict the configuration of the continents during periods of apparent vicariant and dispersal events in ostariophysan fishes. The plot at the right is a comparison of node age estimates from this study and previous ones. Note that our node age estimates for several order-level lineages date to just after the K-Pg Mass Extinction, and far after the fragmentation of Pangaea. Illustrations are by Julia Johnson (lifesciencestudios.com).

Across our time-calibrated phylogeny, all ostariophysan orders contain radiations restricted to individual continents that coincide with the Cretaceous-Paleogene Mass Extinction 66.02 million years ago. *Cithariniformes*, *Gymnotiformes*, African lineages of *Characiformes*, shellears (*Kneriidae*)*, Siluroidei* and a subclade of loricarioid catfishes, two of the three major lineages of *Characiformes* endemic to South America, and all cypriniforms beside *Gyrinocheilus* appear within six million years of the Cretaceous-Paleogene boundary (Figure 1, Figure 2). Except for siluroid catfishes, the initial radiations of these lineages are all restricted to individual continents (Figure 1). These results reconcile the much older, Cretaceous ages previously estimated for most families of *Characiformes* [13,31,32] with the relaxed-molecular clock ages using genomic data for other ostariophysan lineages, such as loricariid catfishes [44], and suggest that rather than representing an ancient set of lineages surviving from the Mesozoic [32], African lineages of *Characiformes* originated from a single Paleogene dispersal from South America. Thus, our time-calibrated phylogeny demonstrates a global signature of lineage turnover in ostariophysan fishes occurring directly after the most recent mass extinction event in Earth history.

Our results also clarify the origins of freshwater ostariophysans and their deep-sea relatives. We resolve only two freshwater invasions in *Ostariophysi*: one in the common ancestor of shellears (*Kneriidae*) and Hingemouth *Phractolaemus ansorgii* (Figure 3), and the other in crown *Otophysi*. These freshwater invasions and the deep-sea radiation of alepocephaliform fishes are alternately associated with the development or degeneration of structures associated with enhanced hearing capacity. Otophysans share the Weberian apparatus, a specialized structure of modified anterior vertebrae and ribs that functionally integrates the swimbladder and inner ear to enhance auditory capacity [28,94,95]. A somewhat analogous condition also occurs in *Clupeiformes*, where the swim bladder capsule anteriorly expands to contact the inner ear via gas-filled bullae [43,96–98]. In contrast, alepocephaliforms have secondarily lost the a swim bladder [25,26]. Other pelagic deep sea fishes, including deep-sea anglerfishes (56,57) also lack a swim bladder, a modification that is associated with the energetic cost of buoyancy regulation at depth [101]. Thus, the loss of the swim bladder in slickheads contrasts with its co-option into the auditory systems of both clupeiforms and ostariophysans and suggests that, rather than representing the retention of the deep marine ecology perhaps ancestral for ray-finned fishes [102,103], alepocephaliforms secondarily invaded the deep sea, where they became some of the deepest-venturing and largest predatory bathypelagic vertebrates [104].

**Figure 3.**
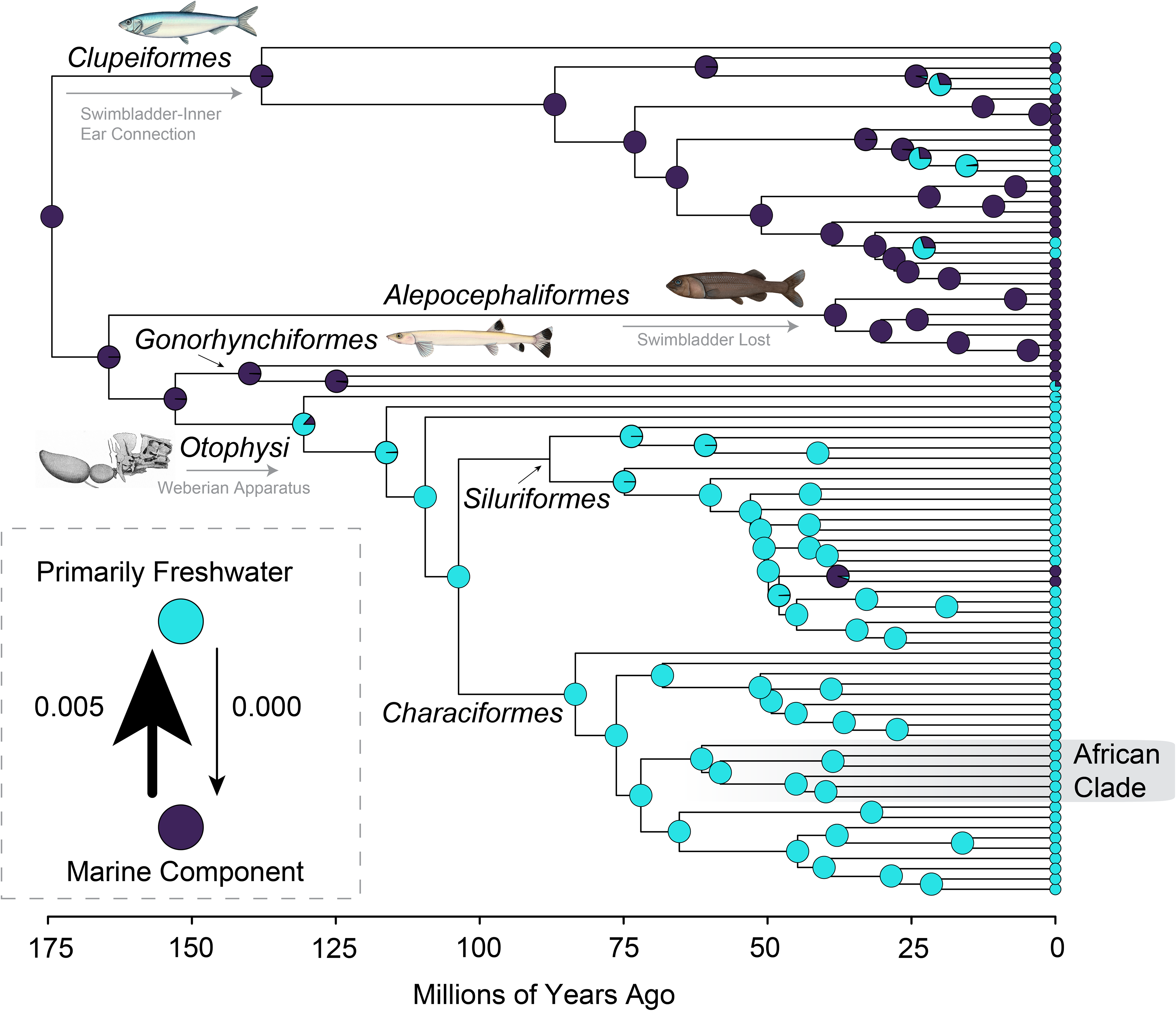
Auditory Systems and Related Structures Track Freshwater and Deep Sea Invasions. Ancestral state reconstruction of habitat coded as a binary trait (freshwater vs. marine component) overlayed on a simplified version of the phylogeny presented in Figure 1. Pie charts indicate the probability of particular states at nodes, and dots at tips denote tip states (pie charts for the collapsed *Cypriniformes* and *Kneriidae* MRCA nodes are shown). Inset shows the transition matrix with arrows denoting transitions between states scaled to match the values of the calculated transition rates. Also annotated with grey text and arrows are gains and losses of auditory structures and associated components in lineages of otocephalans. Illustrations are by Julia Johnson (lifesciencestudios.com).

## Discussion

Here, we show that the evolutionary history of freshwater fish diversity in the mega-radiation *Ostariophysi* [5,6,16,24,44–46,87,94,105–107] is reflective of a pattern of Cretaceous divergences among major lineages such as catfishes, minnows, knifefishes and tetras, followed by sudden lineage origination over the last 65 million years of Earth history (Figure 1). This involved a single transition to freshwater, with rare subsequent dispersals across shallow seas. Many studies have estimated times among ostariophysan lineages endemic to continents that overlap with the fragmentation of Pangaea, implying that ostariophysan diversity is explained by the steady accumulation of lineage diversity through deep time [8,10,11,13,31,32,35,108,109]. This perspective of freshwater fish diversification supports the classic ‘museum’ hypothesis for why the tropics are so species-rich [6,13,110]. However, recent studies that incorporate the most well-characterized fossils of ostariophysans and related lineages as tips [24] or estimate the timescale of ray-finned fish evolution more broadly without densely sampling ostariophysan diversity [18,21,22,27,33,69] estimate similar younger ages for the lineages of *Ostariophysi* that they include (Figure 2). Unlike previous studies that have inferred much older ages, we employ the same fossil calibrations but enhance our approach by extensively sampling the clades most closely related to *Ostariophysi* (Figure 2), thereby avoiding the need to specify a wide prior on the root of this lineage due to its uncertain age. By resolving *Alepocephaliformes* as the sister lineage of *Ostariophysi*, we can more accurately determine the time frame for the origin of the more than 11,700 species found within this clade across Temperate, Neotropical, and Afrotropical regions.

The most striking pattern in our time-calibrated phylogeny is the near-simultaneous origin of many species-rich lineages of ostariophysans, including the taxonomic orders and suborders *Cithariniformes*, *Gymnotiformes*, and *Siluroidei*, within 7 million years of the Cretaceous-Paleogene boundary (Figure 1). A direct reading of the fossil record suggests that freshwater ecosystems were less impacted by the extinction event than terrestrial or marine environments [36–40,111–113]. Consequently, many relaxed-molecular clock analyses that have focused on particular ostariophysan lineages tend to estimate Mesozoic ages for the oldest nodes in the ostariophysan phylogeny, a conclusion that molecular biologists have generally accepted [8,10,11,13,31,32,35]. In contrast, our analyses indicate that freshwater fish biodiversity underwent significant turnover around the K-Pg boundary. This is not entirely surprising with respect to the ostariophysan fossil record, as no unambiguous fossils of crown *Characiformes, Cithariniformes*, *Cypriniformes*, *Kneriidae*, or *Gymnotiformes* are known from the Mesozoic [12,13,24,29,32,44], but does differ considerably from some species-rich time trees of ostariophysan lineages [8,13,31,32,108,109].

Nearly all Mesozoic fishes known from mostly complete body fossils with affinities to ostariophysans have uncertain phylogenetic relationships among the living orders [24,30,114], and fossil occurrences from the Mesozoic used to calibrate phylogenies of living species are comprised of isolated teeth and bones that cannot be confidently referred to subordinal-level lineages, or worse, represent entirely unrelated clades. For example, isolated teeth have been used to calibrate the common ancestors of families of characiform fishes in studies that recover Triassic-Jurassic ages for the deepest divergences in *Ostariophysi* [13], but the placement of these fossils has not been justified on the basis of phylogenetic analyses of morphological characters. These occurrences are also problematic because of growing evidence for the convergent evolution of characiform-like teeth in extinct fish clades [115,116]. The discovery of Jurassic, Cretaceous, and Paleogene species in †*Pycnodontiformes* that possess characiform-like multicuspid shearing dentition [115,116] warrants reexamination of the earliest putative characiform fossils. Combined with the use of mitochondrial DNA in previous studies [8,109,120], which can artificially inflate divergence times due to the prevalence of sequence saturation across the mitochondrial genome [121], the placement of these isolated fossils as calibrations deep within the major ostariophysan crown clades has likely led to the gross overestimation of divergence times across the largest freshwater vertebrate radiations across studies that deploy them [8,13,31,32,109]. The hypothesis of the timescale of ostariophysan evolution that we present, which is based on nearly as many fossil calibrations as the most fossil-rich tip-dated phylogenies of the clade presented in the literature [24], suggests that incongruences between the ages of ostariophysan clades estimated by molecular phylogenies or a literal reading of the fossil record are due to a reliance on isolated, poorly characterized material, a problem that is slowly being rectified as more complete ostariophysans are described from the Mesozoic and early Cenozoic [24,30,118].

The time-calibrated phylogeny and historical biogeographic reconstruction of *Ostariophysi* that we present align with a recently proposed model of regionalized extinction in freshwater ecosystems at the close of the Cretaceous where impacts were varied across local communities and clades [36,122]. Our analyses support the Neotropics as the origin of nearly of the lineages that originate around the K-Pg extinction (Figure 1). Several other freshwater lineages from the southern hemisphere, such as *Mordacia* and *Geotria* lampreys, [123,124], three lineages of lungfishes [125–129], and polypterids [130] also persisted through the mass extinction event. Similar to what has been proposed for mammals [122], it is plausible that the initial diversification of ostariophysans during the Mesozoic enabled them to rapidly exploit the ecological niches left vacant by the casualties of the asteroid impact in the Yucatan. Nonetheless, our findings suggest that the impact of the K-Pg extinction’s role in shaping present-day freshwater biodiversity has been underestimated.

Our results also clarify the timing and number of freshwater invasions and subsequent dispersals in otocephalan fishes. First, we confirm that *Alepocephaliformes* represent a recent, secondary invasion of the bathypelagic and abyssopelagic zones of the world’s oceans, rather than reflecting a possible deep marine ecology that has been considered ancestral for many ray-finned fishes [102]. We infer only two freshwater invasions by ostariophysans, which is concordant with the fossil record of early crown ostariophysans from marginal and shallow marine habitats [24,30]. This history of freshwater and deep marine invasions is tracked by the evolution of enhanced hearing capabilities in otophysans and clupeiforms that exclusively occupy freshwater, brackish, and shallow marine environments (Figure 4). In contrast, *Alepocephaliformes* secondarily lost the swim bladder, which plays a critical functional role in the specialized auditory structures of both *Clupeiformes* and *Ostariophysi* [28,43,94–98]. Thus, the same anatomical features have been secondarily modified or lost in association with major ecological transitions across ostariophysans and their closest relatives.

The historical biogeography of major lineages of catfishes and tetras that we reconstruct suggests that several oceanic dispersals have occurred over the last 65 million years. In catfishes, this is less surprising since several lineages, such as *Ariidae* and *Plotosidae*, include numerous marine species [131–133] and exhibit complicated histories of freshwater colonizations [131,134]. The divergence of African and South American characiforms in the Late Cretaceous is more unexpected, implying that this exclusively freshwater clade dispersed across the Atlantic and did not originate from a Mesozoic radiation initiated by the fragmentation of Gondwana [32]. Due to their low salinity tolerance [135], hypotheses explaining trans-Atlantic dispersals in *Characiformes* have relied on ephemeral land connections, reduced-salinity corridors, or chance dispersals via rare events like rafting [136]. Contrary to one recent study [69], the recent arrival of characiforms to Africa cannot be explained by the persistence and subsequent extinction of South American lineages on the African continent; this explanation is not compatible with the nested position of African lineages among South American clades or the clade ages we estimate.

A Paleogene dispersal of tetras, pikes, and tigerfishes to Africa might be explained by ancestral physiological adaptations. The earliest divergence in the sister lineage to African characiforms consists of *Tarumania walkerae* and *Erythrinidae*, which are notable for the ability to facultatively breathe air [31,137,138]. It is conceivable that saltwater tolerance may have been independently lost in African tetras and the two or three major lineages of South American characiforms over the 70 million year history of the *Characiformes*. The history of *Characiformes* might reflect of ancient physiological capacities that have subsequently been lost through extinction and evolution in modern lineages with a long history of exclusively freshwater inhabitation.

Science are increasingly reaching a consensus on the phylogenetic relationships and evolutionary timescales of branches on the Tree of Life that have historically been difficult to resolve [144–149]. Our inference of the relationships of *Ostariophysi* using genomic data allows us to more confidently estimate the timescale over which the most diverse freshwater vertebrate lineages on the planet [6,13,21,44,46,110] accumulated their diversity. By demonstrating that the most species-rich lineages of ostariophysans all appeared shortly after the Cretaceous-Paleogene mass extinction, our results suggest that the impact the most recent of Earth’s big five [150,151] mass extinctions, on freshwater ecosystems [36–40,112] has been significantly underestimated. The hyperdiverse freshwater fish faunas of the tropics accumulated in a world recovering from the Chicxulub impact.

## Competing Interests

The authors declare that they have no competing interests.

## Data Availability

All data from this study is available in the Supplementary Information associated with this article or the associated Yale Dataverse Repository (https://doi.org/10.60600/YU/8AWZYX). New sequences will be uploaded to NCBI GenBank sequence archive under the BioProject: XXX.

## Notes

### Competing Interest Statement

The authors have declared no competing interest.

### Funding

- National Institute of General Medical Sciences: 5T32GM148332-03
- U.S. National Science Foundation: DEB-2508461
- Yale Peabody Museum: Bingham Oceanographic Fund
- Society of Systematic Biologists: Graduate Student Research Award
- Fundação de Amparo à Pesquisa do Estado de São Paulo: 2024/19358-7, 2020/13433-6
- CNPq: 306054/2006-0
- CNPq: 441128/2020-3

