## Supplementary Information for "The largest radiations of freshwater fishes initiated at the Cretaceous-Paleogene boundary"

**Supplement To: Genomes and auditory structures track the assembly of freshwater fish biodiversity**

**Fossil Calibration List.**

†*Ichthyemidion vidali*

**Justification of Placement:** †*Ichthyemidion vidali* calibrates the MRCA of *Oseanacephala* (specifiers: *Megalops atlanticus*, *Scleropages formosus*) in our node-dating analysis. The placement of †*I. vidali* as a member of total group *Elopomorpha* is supported by Bayesian and parsimony analyses, including the morphological + molecular dataset of Dornburg et al. (1), the 74 character, 27 taxon dataset of Figueiredo et al. (2), and the 75 character, 30 taxon dataset of Hernández-Guerrero et al. (3). Given the considerable uncertainty surrounding the ages and phylogenetic positions of Late Jurassic to Early Cretaceous members of *Elopomorpha* and its constituent clades (1, 2, 4, 5), this is a conservative fossil calibration.
**Stratigraphic Horizon**: El Montsec, Lérida Province, Spain; Berriasian–Barremian (1). We use an age of 125.0 Ma, which is the traditional upper bound of the Barremian (6).
**Fossil tip age**: 125.0 Ma. The node calibration was created such that 97.5% of the probability distribution fell before 125.0 Ma.

†*Tischlingerichthys viohli*

**Justification of Placement:** †*Tischlingerichthys viohli* calibrates the MRCA of *Otocephala* (specifiers: *Clupea harengus*, *Danio rerio*) in our node-dating analysis. The placement of †*T. viohli* in crown *Otocephala* is supported by parsimony analyses of morphological characters, including the 74 character, 27 taxon dataset of Figueiredo et al. (2) and the 75 character, 31 taxon matrix of L-Recinos et al. (7). For the purposes of this justification, we rely on the phylogeny presented in figure 15 of L-Recinos et al. (7).
**Stratigraphic Horizon**: Mühlheim, Bavaria, Germany; Mörnsheim Formation, lower Tithonian (4). We use an age of 151.5 Ma, which is the maximum upper bound on the age of the Tithonian (6).
**Fossil tip age**: 151.5 Ma. The node calibration was created such that 97.5% of the probability distribution fell before 151.5 Ma.

†*Pseudoellimma gallae*

**Justification of Placement:** †*Pseudoellimma gallae* calibrates the MRCA of *Clupeiformes* (specifiers: *Denticeps clupeoides*, *Clupea harengus*) in our node-dating analysis. The placement of †*P. gallae* as a member of *Clupeiformes* with affinities to *Clupeoidei* is supported by: presence of accessory anterior foramen of temporal canal, postorbital branch of supraorbital canal on frontal not visible in lateral view; skull roof bone texture striated; ural centrum I reduced (8).
**Stratigraphic Horizon**: Atol Quarry, São Miguel dos Campos, Alagoas State, Brazil; Coqueiro Seco Formation, Barremian Stage of the Cretaceous (8). We use an age of 125.0 Ma, which is the traditional upper bound of the Barremian (6) following previous studies (9).
**Fossil tip age**: 125.0 Ma. The node calibration was created such that 97.5% of the probability distribution fell before 125.0 Ma.

†*Clupeopsis straeleni*

**Justification of Placement:** †*Clupeopsis straeleni* calibrates the MRCA of *Engraulidae* and *Spratelloididae* (specifiers: *Anchoviella jamesi*, *Jenkinsia* sp.) in our node-dating analysis. The placement of †*C. straeleni* as a member of Pan-*Engraulidae* is supported by: posteriorly inclined suspensorium, mesethmoid projects anterior to vomers; large proportion of metapterygoid placed anterodorsal to quadrate; hyomandibula ventral process reaches posterior end of quadrate; maxilla straight; reduced coronoid process; expanded lateral laminar process of hyomandibula overlaps metapterygoid (10).
**Stratigraphic Horizon**: Dubois clay pit, Chièvres, Hainaut, Belgium; Lower Orchies Clay Member, Ypresian, Eocene, 54.40 to 54.05 Ma (10).
**Fossil tip age**: 54.05 Ma. The node calibration was created such that 97.5% of the probability distribution fell before 54.05 Ma.

†*Rubiesichthys gregalis*
**Justification of Placement:** †*Rubiesichthys gregalis* calibrates the MRCA of *Chanos chanos* and *Phractolaemus ansorgii* in our node-dating analyses. The placement of †*R. gregalis* in *Ostariophysi* in *Gonorhynchiformes* as a member of Pan-*Chanos* is supported by Bayesian and parsimony analyses of morphological characters, including the 94 character, 22 taxon matrix of Grande and Poyato-Ariza (11), the 106 character, 14 taxon matrix of Ribiero et al. (12), the 130 character, 22 taxon matrix of Grande et al. (13) (including as modified by Amaral et al. (14)), and the 128 character, 20 taxon matrix used with molecular data by Near et al. (15).
**Stratigraphic Horizon**: El Montsec and Las Hoyas, Lérida and Cuenca Provinces, Spain (16, 17); Berriasian–Barremian (1). El Montsec is Barriasian through lower Barremian, whereas Las Hoyas is Barremian (18, 19). We use an age of 121.4 Ma, which is the upper bound of the Barremian (6).
**Fossil tip age**: 121.4 Ma. The node calibration was created such that 97.5% of the probability distribution fell before 121.4 Ma.

†*Mahengichthys singidaensis*
**Justification of Placement:** †*Mahengichthys singidaensis* calibrates the MRCA of *Kneriidae* (specifiers: *Kneria ruaha*, *Grasseichthys gabonensis*) in our node-dating analysis. The placement of †*M. singidaensis* in *Kneriidae* as a member of Pan-*Kneria* is supported by Bayesian, maximum likelihood, and parsimony analyses of morphological characters, including the 128 character, 23 taxon matrix of Davis et al. (20).
**Stratigraphic Horizon**: Mahenge locality, near Mwaru village, 65 km west of Singida, Tanzania; Mahenge Formation, Lutetian, Eocene, 46 to 45 Ma (20).
**Fossil tip age**: 45.0 Ma. The node calibration was created such that 97.5% of the probability distribution fell before 45.0 Ma.

†*Santanichthys diasii*
**Justification of Placement:** †*Santanichthys diasii* calibrates the MRCA of *Otophysi* (specifiers: *Corydoras julii* (= *Hoplisoma julii*) , *Danio rerio*) in our node-dating analyses. The placement of †*S. diasii* in *Otophysi* is supported by the following characters: basisphenoid absent, dermopalatine absent, expanded dorsomedial portion of anterior neural arches, expansion of the anterior supraneural, modification of the first neural arch into the scaphium, modification of the second neural arch into the intercalarium, first four centra shortened, and modification of the third centrum rib parapophysis into the tripus (21).

**Stratigraphic Horizon**: Araripe Basin, northeastern Brazil; Santana Group, Albian Stage of the Early Cretaceous (21). Following previous studies, we use an age of 112.5 Ma (22, 23).
**Fossil tip age**: 112.5 Ma. The node calibration was created such that 97.5% of the probability distribution fell before 112.5 Ma.

†*Wilsonium brevipinne*
**Justification of Placement:** †*Wilsonium brevipinne* calibrates the MRCA of *Catostomidae* and *Danioidae* (specifiers: *Catostomus latipinnis*, *Danio rerio*) in our node-dating analyses. The placement of †*W. brevipinne* in *Cypriniformes* as a member of Pan-*Catostomidae* is supported by parsimony analysis of morphological characters, including the 83 character, 71 taxon matrix of Liu et al (24). For the purposes of this justification, we rely on the phylogeny presented in figure 6 of Liu et al (24). Phylogenetically optimized apomorphies that unite †*W. brevipinne* with Pan-*Catostomidae* are: pharyngeal toothplate falcate (24).
**Stratigraphic Horizon**: North Fork of the Similkameen River, Pleasant Valley, British Columbia, Canada; Allenby Formation, Eocene (24). Following Bagley et al. (25), we use an age of 48.88 Ma for this taxon.
**Equivalent Fossil Calibrations:** Various pan-catostomids, see Bagley et al. (25).
**Fossil tip age**: 48.88 Ma. The node calibration was created such that 97.5% of the probability distribution fell before 48.88 Ma.

†*Cobitis nanningensis*
**Justification of Placement:** †*Cobitis nanningensis* calibrates the MRCA of *Cobitis sinensis* and *Misgurnus anguillicaudatus* in our node-dating analysis. The placement of †*C. nanningensis* as a member of Pan-*Cobitis* is supported by the presence and shape of the suborbital spine (26).
**Stratigraphic Horizon**: Nanning Basin, Guangxi Zhuang Autonomous Region, southern China; Lower Yongning Formation, early to middle Oligocene (Rupelian) (26), 33.9 to 27.3 Ma (6).
**Fossil tip age**: 27.3 Ma. The node calibration was created such that 97.5% of the probability distribution fell before 27.3 Ma.

†*Eoprocypris maomingensis*
**Justification of Placement:** †*Eoprocypris maomingensis* calibrates the MRCA of *Cyprininae* and *Smiliogastrinae* (specifiers: *Enteromius treurensis*, *Cyprinus carpio*) in our node dating analysis. The placement of †*E. maomingensis* as a member of Pan-*Cyprininae* is supported by: presence of five branched anal fin rays, serrated last unbranched dorsal and anal fin rays (27). †*E. maomingensis* is also united with *Procypris* among living cyprinines based on: spoon-like grinding surface on pharyngeal teeth and anterior and posterior processes of pharyngeal toothplate equal in length (27).
**Stratigraphic Horizon**: Jintang, Maoming County, Guangdong Province, China; Youganwo Formation, Chron 18 (27), which extends from 41.03 to ~38 Ma (6).
**Fossil tip age**: 38.0 Ma. The node calibration was created such that 97.5% of the probability distribution fell before 38.0 Ma.

†*Iquius nipponicus*
**Justification of Placement:** †*Iquius nipponicus* calibrates the MRCA of *Xenocyprididae* (specifiers: *Megalobrama amblycephala*, *Ctenopharyngodon idella*) in our node-dating analysis. The placement of †*Iquius nipponicus* as a member of *Xenocyprididae* on the stem of

*Xenocyprididae* is supported by phylogenetic analysis of morphological characters, including the 24 character, 14 taxon matrix of Yabumoto and Sakamoto (28).
**Stratigraphic Horizon**: Diatomite at Hachiman, Ashibe, Iki Island Nagasaki Prefecture, Japan; Chojabaru Formation, Iki Group, Middle Miocene, 15.3 Ma (28).
**Fossil tip age**: 15.3 Ma. The node calibration was created such that 97.5% of the probability distribution fell before 15.3 Ma.

†*Hydrocynus* sp.
**Justification of Placement:** †*Hydrocynus* sp. calibrates the MRCA of *Alestes baremoze* and *Brachypetersius altus* our node-dating analysis. The placement of †*Hydrocynus* sp. in *Alestidae* as a member of *Hydrocynus*, the living sister taxon (29) to *Alestes*, based on the following combination of features: pointed, unicuspidate, labiolingually compressed teeth with a slightly concave lingual and convex labial face (30).
**Stratigraphic Horizon**: Dur At-Talah, Sirt Basin, Libya; Bioturbated Unit, Chron 18n.1n, 39 to 38 Ma (30).
**Fossil tip age**: 38.0 Ma. The node calibration was created such that 97.5% of the probability distribution fell before 38.0 Ma.

†*Paleotetra entrecorregos*
**Justification of Placement:** †*Paleotetra entrecorregos* calibrates the MRCA of *Characidae* and *Acestrorhamphidae* (specifiers: *Charax condei*, *Oxybrycon parvulus*) in our node-dating analysis. The placement of †*Paleotetra entrecorregos* as a member of Pan-*Characidae* is supported by the following combination of features: long anal fin, toothed maxilla, 10-13 rays in the dorsal fin, 12 dorsal, nine ventral procurrent caudal fin rays with the anteriormost ventral procurrent caudal fin ray plate-like (31).
**Stratigraphic Horizon**: Ravines of the Entre-Córregos stream, near Aiuruoca, Minas Gerais State, Brazil; Entre-Córregos Formation, Eocene-Oligocene, 33.9 Ma (31).
**Fossil tip age**: 33.9 Ma. The node calibration was created such that 97.5% of the probability distribution fell before 33.9 Ma.

†*Bagridae* indet. BQ-2
**Justification of Placement:** †*Bagridae* indet. calibrates the MRCA of *Bagridae* and *Schilbeidae* (specifiers: *Bagrus docmak*, *Eutropiichthys vacha*) in our node-dating analysis. The placement of †*Bagridae* indet. material from the BQ-2 locality a member of Pan-*Bagridae* is supported by: very thin anteroposterior width of vertebral centra in lateral view, pectoral fin spines lack tubercles on the anterior surface, ventral and axial processes of pectoral fin spines strongly developed (32).
**Stratigraphic Horizon**: Birket Qarun locality 2, Fayum Depression, Egypt; Umm Rigl Member, Birket Qarun Formation, lower Priabonian, Eocene, ~37 Ma (32).
**Fossil tip age**: 37.0 Ma. The node calibration was created such that 97.5% of the probability distribution fell before 37.0 Ma.

†*Qarmoutus hitanensis*
**Justification of Placement:** †*Qarmoutus hitanensis* calibrates the MRCA of *Ariidae* and *Plotosidae* (specifiers: *Arius maculatus*, *Plotosus canius*) in our node-dating analysis. The

placement of †*Q. hitanensis* as a member of Pan-*Ariidae* is supported by phylogenetic analysis of morphological characters, including the 230 character, 94 taxon matrix of El-Sayed et al. (33).
**Stratigraphic Horizon**: Wadi El-Hitan, Fayum Depression, Egypt; Birket Qarun Formation, Priabonian, Eocene (33), 37.71 to 33.9 Ma (6). Note that indeterminate isolated bones possibly assignable to *Ariidae* are known from the BQ-2 locality, which is early Priabonian, 37.71 Ma (32). However, we conservatively use †*Qarmoutus hitanensis* as the calibration given that it is known from more complete material.
**Fossil tip age**: 33.9 Ma. The node calibration was created such that 97.5% of the probability distribution fell before 33.9 Ma.

### III. References.

6. F. M. Gradstein, J. G. Ogg, M. Schmitz, G. Ogg, *The Geologic Time Scale 2020* (Elsevier Science, 2021).

21. A. Filleul, J. G. Maisey, Redescription of Santanichthys diasii (Otophysi, Characiformes) from the Albian of the Santana Formation and comments on its implications for otophysan relationships. American Museum novitates ; no. 3455. (2004).

**Figure S1. Maximum likelihood phylogenies of *Otocephala* based on the complete dataset.** The phylogeny is from the analysis where the concatenated sequence data were treated as a single partition.

**Figure S2. Maximum likelihood phylogenies of *Otocephala* based on the complete dataset.** The phylogeny is from the analysis where the concatenated sequence data were treated as multiple partitions.

**Figure S3. ASTRAL-III multispecies coalescent phylogeny of *Otocephala* based on the complete dataset.**

**Figure S4. Maximum likelihood phylogeny of *Otocephala* based on the reduced dataset.** The phylogeny is from the analysis where the concatenated sequence data were treated as a single partition.

**Figure S5. Maximum likelihood phylogenies of *Otocephala* based on the reduced dataset.** The phylogeny is from the analysis where the concatenated sequence data were treated as multiple partitions.

**Figure S6. ASTRAL-III multispecies coalescent phylogeny of *Otocephala* based on the reduced dataset.**

**Figure S7. Phylogenomic Resolution of the Ostariophysan Sister Lineage.** Simplified phylogenies from the single-partition concatenated analyses of the complete and subsampled datasets showing the relationships of the major otocephalan lineages. Numbers at nodes and branches of major lineages indicate the bootstrap (left), gene (middle), and site (right) concordance factors supporting the resolution of the associate nodes or the monophyly of the associated order-level lineage. Illustrations are by Julia Johnson (lifesciencestudios.com).

**Figure S8. Comparison of dataset sizes.** Barchart compares the sequence dataset size of our study and previous ones dealing with otocephalan relationships.

**Figure S9. Gene and site concordance factors.** Plots show the relationships between gene and site concordance factors, ultrafast bootstrap supports, and log-transformed branch lengths calculated for the complete (top row) and reduced (bottom row) datasets.

**Figure S10. Fossil Calibration Scheme.** (A) Simplified version of our time-calibrated phylogeny showing placement of fossil calibrations along the tree. (B) Comparisons of prior and posterior age distributions for calibrated nodes.

**Figure S11. Ancestral state reconstruction of habitat preference.** Ancestral state reconstruction of habitat preference where habitat is treated as a polymorphic multistate character. Pie charts indicate the probability of particular states at nodes, and dots at tips denote tip states (pie charts for the collapsed Cypriniformes and Kneriidae MRCA nodes are shown). Inset shows the transition matrix with arrows denoting transitions between states scaled to match the values of the calculated transition rates. Illustrations are by Julia Johnson (lifesciencestudios.com).

**Figure S12. Simplified historical biogeographic reconstruction.** Biogeographic reconstruction from BioGeoBEARS resulting from mapping using the best-fit model (BAYAREALIKE) on the three-area scheme. Pie charts at nodes indicate probabilities for the node occupying particular areas in the historical biogeographic reconstruction conducted in BioGeoBEARS. Illustrations are by Julia Johnson (lifesciencestudios.com).

**Table S1. List of UCE sources.**

| **Species** | **Tip Identifier** | **Source** |
|---|---|---|
| *Osteoglossum bicirrhosum* | osteoglossum_bicirrhosum_JBIPBQ01 | GCA_047301455 |
| *Scleropages formosus* | scleropages_formosus_CAAHFQ01 | GCA_900964775.1 |
| *Albula goreensis* | albula_goreensis_JAERUA01 | GCA_022829145.1 |
| *Megalops atlanticus* | megalops_atlanticus_JAFDVH01 | GCA_019176425.1 |
| *Argentina brasiliensis* | argentina_brasiliensis_102017 | New |
| *Borostomias antarcticus* | teleostei_borostomias_antarcticus_CATLJS01 | GCA_949987555.1 |
| *Denticeps clupeoides* | denticeps_clupeoides_caadhs02 | GCA_900700375.2 |
| *Jenkinsia sp.* | Jenkinsia_sp_2465 | New |
| *Lycengraulis gossidens* | lycengraulis_gossidens_48719 | New |
| *Amazonsprattus scintilla* | amazonsprattus_scintilla_57222 | New |
| *Anchoviella jamesi* | anchoviella_jamesi_108920 | New |
| *Sprattus sprattus* | sprattus_sprattus_cauoqg01 | GCA_963457725.1 |
| *Clupea harengus* | clupea_harengus_caadhv01 | GCA_900700415.2 |
| *Clupea pallasii* | clupea_pallasii_jbpkrg01 | GCA_051176545.1 |
| *Chirocentrodon bleekerianus* | chirocentrodon_bleekerianus_48602 | New |
| *Ilisha elongata* | ilisha_elongata_jblrxz01 | GCA_048126385.1 |
| *Pellona flavipinnis* | pellona_flavipinnis_86425 | New |
| *Ilisha amazonica* | ilisha_amazonica_106163 | New |
| *Pristigaster cayana* | pristigaster_cayana_57107 | New |
| *Alosa pseudoharengus* | alosa_pseudoharengus_jbggwu01 | GCA_046254945.1 |
| *Alosa sapidissima* | alosa_sapidissima_18508 | New |
| *Alosa sapidissima* | alosa_sapidissima_jahdtn01 | GCA_018492685.1 |
| *Sardina pilchardus* | sardina_pilchardus_cawkyn01 | GCA_963854185.1 |
| *Sardinops sagax* | sardinops_sagax_18521 | New |
| *Sardinops sagax* | sardinops_sagax_jbiekb01 | GCA_044707015.1 |
| *Herklotsichthys quadrimaculatus* | herklotsichthys_quadrimaculatus_jbnwuu01 | GCA_051253255.1 |
| *Konosirus punctatus* | konosirus_punctatus_15924 | New |
| *Rhinosardinia amazonica* | rhinosardinia_amazonica_46985 | New |
| *Sardinella tawilis* | sardinella_tawilis_jascqy01 | GCA_030264315.1 |
| *Dorosoma cepedianum* | Dorosoma_cepedianum_t14313 | New |
| *Harengula clupeola* | harengula_clupeola_46934 | New |

| | | |
|---|---|---|
| *Sardinella aurita* | sardinella_brasiliensis_82073 | New |
| *Holtbyrnia latifrons* | holtbyrnia_latifrons_sio18516 | New |
| *Mirorictus taningi* | mirorictus_taningi_sio18518 | New |
| *Talismania bifurcata* | talismania_bifurcata_sio18522 | New |
| *Xenodermichthys copei* | xenodermichthys_copei_102283 | New |
| *Conocara macropterum* | conocora_macropterum_102239 | New |
| *Alepocephalus tenebrosus* | alepocephalus_tenebrosus_sio18506 | New |
| *Alepocephalus agassizii* | alepocephalus_agassizii_jbnttr01 | GCA_051994585.1 |
| *Gonorhynchus greyi* | gonorhynchos_gery_ams18659 | New |
| *Chanos chanos* | chanos_chanos_sio18514 | New |
| *Chanos chanos* | chanos_chanos_cabivn01 | GCA_902362185.1 |
| *Phractolaemus ansorgii* | Phractolaemus_ansorgii_YFTC_19988 | New |
| *Kneria paucisquamata* | Kneria_paucisquamata_YFTC_18542 | New |
| *Kneria ruaha* | Kneria_ruaha_YFTC_14489 | New |
| *Cromeria nilotica* | cromeria_nilotica_221901 | New |
| *Grasseichthys gabonensis* | grasseichthys_gabonensis_205322 | New |
| *Parakneria abbreviata* | parakneria_abbreviata_211882 | New |
| *Parakneria spekii* | Parakneria_spekii_YFTC_14483 | New |
| *Parakneria ladigesi* | parakneria_ladigesi_255286 | New |
| *Parakneria vilhenae* | parakneria_vilhenae_249152 | New |
| *Gyrinocheilus aymonieri* | gyrinocheilus_aymonieri_jbpgvv01 | GCA_051167395.1 |
| *Catostomus latipinnis* | catostomus_latipinnis_jazhbr01 | GCA_036785435.1 |
| *Botia almorhae* | botia_almorhae_jazgtj01 | GCA_036877575.1 |
| *Cobitis sinensis* | cobitis_sinensis_jbnunk01 | GCA_051993705.1 |
| *Misgurnus anguillicaudatus* | misgurnus_anguillicaudatus_jaldxj02 | GCA_027580225.2 |
| *Barbatula barbatula* | barbatula_barbatula_jaxofq01 | GCA_947034865.1 |
| *Triplophysa tibetana* | triplophysa_tibetana_soyy01 | GCA_008369825.1 |
| *Beaufortia kweichowensis* | beaufortia_kweichowensis_jaggec01 | GCA_019155185.1 |
| *Lepturichthys fimbriata* | lepturichthys_fimbriata_jbpgzu01 | GCA_051168995.1 |
| *Ctenopharyngodon idella* | ctenopharyngodon_idella_jaiklg01 | GCA_019924925.1 |
| *Megalobrama amblycephala* | megalobrama_amblycephala_jagtwi01 | GCA_018812025.1 |
| *Gobio gobio* | gobio_gobio_cashtd01 | GCA_949357685.1 |
| *Leuciscus idus* | leuciscus_idus_jajavr01 | GCA_021554675.1 |
| *Squalius caroltertii* | squalius_carolitertii_japmod01 | GCA_029875075.1 |
| *Phoxinus dragarum* | phoxinus_dragarum_jarpmj01 | GCA_033366665.1 |
| *Gila robusta* | gila_robusta_javalu01 | GCA_030770095.1 |
| *Rhinichthys osculus* | rhinichthys_osculus_jarwkj01 | GCA_029959095.1 |
| *Danio rerio* | danio_rerio_jbmgra01 | GCA_049306965.1 |
| *Garra ornata* | garra_ornata_jbmfzy01 | GCA_049181675.1 |
| *Labeo gonius* | labeo_gonius_npdk01 | GCA_013461565.1 |

| | | |
|---|---|---|
| *Barbus barbus* | barbus_barbus_cakzfl01 | GCA_936440315.1 |
| *Carassius auratus* | carassius_auratus_qpke01 | GCA_003368295.1 |
| *Acrossocheilus wenchowensis* | acrossocheilus_wenchowensis_jajuba01 | GCA_054650785.1 |
| *Enteromius treurensis* | enteromius_treurensis_jaucmu01 | GCA_032275025.1 |
| *Sinocyclocheilus anshuiensis* | sinocyclocheilus_anshuiensis_lave01 | GCA_001515605.1 |
| *Cyprinus carpio* | cyprinus_carpio_jaeoab01 | GCA_018340385.1 |
| *Eigenmannia vincentespelaea* | eigenmannia_vicentespelaea_62040 | Previously_published |
| *Sternopygus macrurus* | sternopygus_macrurus_46840 | Previously_published |
| *Sternarchorhamphus muelleri* | sternarchorhamphus_muelleri_ansp182579 | Previously_published |
| *Apteronotus albifrons* | Aalb | New |
| *Apteronotus albifrons* | apteronotus_albifrons_jbntsm01 | GCA_051992785.1 |
| *Sternarchorhynchus goeldii* | sternarchorhynchus_goeldii_jaoxyf01 | GCA_028565775.1 |
| *Melanosternarchus amaru* | Melanosternarchus_amaru_t17825 | New |
| *Melanosternarchus amaru* | Melanosternarchus_amaru_t17826 | New |
| *Sternarchella calhamazon* | Scal | New |
| *Tenebrosternarchus preto* | tenebrosternarchus_preto_jaoymr01 | GCA_025802075.1 |
| *Gymnotus pantherinus* | GpaLBP37171 | New |
| *Electrophorus voltai* | electrophorus_voltai_jaroks01 | GCA_030913585.1 |
| *Electrophorus electricus* | Eele | New |
| *Electrophorus electricus* | electrophorus_electricus_jabvme01 | GCA_013358815.1 |
| *Rhamphichthys apurensis* | rhamphichthys_apurensis_43111 | New |
| *Steatogenys elegans* | steatogenys_elegans_19728 | New |
| *Steatogenys elegans* | steatogenys_elegans_ansp200421 | New |
| *Brachyhypopomus gauderio* | brachyhypopomus_gauderio_jbqqvn01 | GCA_052324685.1 |
| *Brachyhypopomus brevirostris* | brachyhypopomus_brevirostris_16705 | New |
| *Brachyhypopomus occidentalis* | Bocc_STRI203 | New |
| *Brachyhypopomus occidentalis* | brachyhypopomus_occidentalis_jagxod01 | GCA_020368025.1 |
| *Citharinus congicus* | citharinus_congicus_818014 | Melo et al. (2022) |
| *Citharinus gibbosus* | citharinus_gibbosus_258320 | New |
| *Neolebias ansorgii* | neolebias_ansorgii_230233 | New |
| *Hemidistichodus vaillanti* | hemistischodus_vaillanti_227076 | New |
| *Mesoborus crocodilus* | mesoborus_crocodilus_258349 | New |
| *Eugnathichthys macroterolepis* | eugnatichthys_macroterolepis_270552 | New |
| *Phago boulengeri* | phago_boulengeri_252460 | Melo et al. (2022) |
| *Nannocharax ansorgii* | nannocharax_ansorgii_11111018 | Melo et al. (2022) |
| *Distichodus affinis* | distichodus_affinis_818098 | Melo et al. (2022) |
| *Distichodus hypostomatus* | distichodus_hypostomatus_19540 | Melo et al. (2022) |
| *Paradistichodus dimidiatus* | paradistichodus_dimidiatus_223072 | New |
| *Distichodus atroventralis* | distichodus_atroventralis_jbmaue01 | Genbank |
| *Distichodus sexfasciatus* | distichodus_sexfasciatus_javgvs01 | Genbank |

| | | |
|---|---|---|
| *Trichogenes longipinnis* | trichogenes_longipinnis_3862 | Ochoa et al. (2020) |
| *Callichthys callichthys* | callichthys_callichthys_12299 | New |
| *Astroblepus homodon* | astroblepus_homodon_91157 | New |
| *Scoloplax dolicholophia* | scoloplax_dolicholophia14337 | New |
| *Diplomystes mesembrinus* | diplomystes_mesembrinus5822 | New |
| *Cetopsis arcana* | cetopsis_arcana_64712 | New |
| *Malapterurus electricus* | Malapterurus_electricus_16121 | New |
| *Mochokus niloticus* | Mochokus_niloticus_t15500 | New |
| *Pangasius macronema* | Pangasius_macronema_114085 | New |
| *Clarias batu* | clarias_batu_ansp17140 | New |
| *Kryptoglanis shajii* | kryptoglanis_shajii_ansp3948 | New |
| *Liobagrus reinii* | liobagrus_reinii_ansp14901 | New |
| *Bagrus docmak* | Bagrus_docmak_t18654 | New |
| *Eutropiichthys vacha* | eutropiichthys_vacha_auft3820 | New |
| *Arius maculatus* | Arius_maculatus_t6055 | New |
| *Plotosus canius* | Plotosus_canius_114495 | New |
| *Bunocephalus coracoideus* | Bunocephalus_coracoideus_50615 | New |
| *Auchenipterus nigripinnis* | auchenipterus_nigripinnis_ansp2090 | New |
| *Wertheimeria maculata* | wertheimeria_maculata37522 | New |
| *Conorhynchus conirostris* | conorhynchus_conirostris89907 | New |
| *Lophiosilurus apurensis* | cephalosilurus_apurensis19182 | New |
| *Pimelodus maculatus* | pimelodus_maculatus61290 | New |
| *Characidium etheostoma* | characidium_etheostoma_86571 | Melo et al. (2022) |
| *Boulengerella cuvieri* | boulengerella_cuvieri61652 | Melo et al. (2022) |
| *Brycon amazonicus* | brycon_amazonicus58483 | Melo et al. (2022) |
| *Clupeacharax anchoveoides* | clupeacharax_anchoveoides5046 | Melo et al. (2022) |
| *Gasteropelecus sternicla* | gasteropelecus_sternichla70214 | Melo et al. (2022) |
| *Acestrorhynchus falcatus* | acestrorhynchus_falcatus34081 | Melo et al. (2022) |
| *Charax condei* | charax_condei_74340 | Melo et al. (2024) |
| *Oxybrycon parvulus* | oxybrycon_parvulus_7090 | Melo et al. (2024) |
| *Trochilocharax ornatus* | trochilocharax_ornatus_63103 | Melo et al. (2024) |
| *Hepsetus cuvieri* | hepsetus_cuvieri_249170 | Melo and Stiassny (2024) |
| *Arnoldichthys spilopterus* | arnoldichthys_spilopterus_102152 | Melo and Stiassny (2024) |
| *Lepidarchus adonis* | lepidarchus_adonis_bm108 | Melo and Stiassny (2024) |
| *Brycinus grandisquamis* | brycinus_grandisquamis_227411 | Melo and Stiassny (2024) |
| *Alestes baremoze* | alestes_baremoze_236880 | Melo and Stiassny (2024) |
| *Brachypetersius altus* | brachypetersius_altus_258400 | New |
| *Erythrinus erythrinus* | erythrinus_erythinus_auft6520 | Melo et al. (2022) |

| | | |
|---|---|---|
| *Tarumania walkerae* | tarumania_walkerae_86548 | Melo et al. (2022b) |
| *Argonectes longiceps* | argonectes_longiceps_67384 | New |
| *Piaractus mesopotamicus* | piaractus_mesopotamicus_23804 | New |
| *Serrasalmus maculatus* | serrasalmus_maculatus_22107 | New |
| *Cynodon gibbus* | Cynodon_gibbus_10227_43105 | Melo et al. (2022) |
| *Anostomus anostomus* | anostomus_anostomus_USNM402905 | New |
| *Chilodus gracilis* | chilodus_gracilis_6962_33397 | Melo et al. (2022) |
| *Steindachnerina guentheri* | steindachnerina_guentheri_aum54441_t09661 | New |

**Table S2. Model Fitting for the Ancestral State Reconstruction of Swimbladder Presence.** Bold indicates best-fit model.

| | log(L) | d.f. | AIC | weight |
|---|---|---|---|---|
| ER model | -5.584615 | 1 | 13.16923 | 0.3134888 |
| Irr1 model | -5.836934 | 1 | 13.67387 | 0.2435798 |
| **Irr2 model** | **-5.552225** | **1** | **13.10445** | **0.3238088** |
| ARD model | -5.552225 | 2 | 15.10445 | 0.1191226 |

**Table S3. Model Fitting for the Ancestral State Reconstruction of Extension of the Swimbladder into Neurocranium.** Bold indicates best-fit model.

| | log(L) | d.f. | AIC | weight |
|---|---|---|---|---|
| ER model | -6.558188 | 1 | 15.11638 | 0.2552267 |
| Irr1 model | -6.620996 | 1 | 15.24199 | 0.2396894 |
| **Irr2 model** | **-6.188877** | **1** | **14.37775** | **0.3692459** |
| ARD model | -6.188877 | 2 | 16.37775 | 0.135838 |

**Table S4. Model Fitting for the Ancestral State Reconstruction of the Weberian Apparatus.** Bold indicates best-fit model.

| | log(L) | d.f. | AIC | weight |
|---|---|---|---|---|
| ER model | -7.311255 | 1 | 16.62251 | 0.1887488 |
| **Irr1 model** | **-6.166575** | **1** | **14.33315** | **0.59294248** |
| Irr2 model | -14.281218 | 1 | 30.56244 | 0.00017737 |
| ARD model | -6.166575 | 2 | 16.33315 | 0.21813135 |

**Table S5. Model Fitting for the Historical Biogeographic Reconstruction of the Eight Area Dataset.** Bold indicates best-fit model.

| Model | LnL | numparams | d | e | j | AIC | Weight |
|---|---|---|---|---|---|---|---|
| DEC | -298.2 | 2 | 0.0011 | 0.0002 | 0 | 600.5 | 4.60E-13 |

| | | | | | | | |
|---|---|---|---|---|---|---|---|
| DEC+J | -281.1 | 3 | 0.0009 | 1.00E-12 | 0.012 | 568.3 | 4.50E-06 |
| DIVALIKE | -282.5 | 3 | 0.001 | 1.00E-12 | 0.01 | 571.1 | 1.10E-06 |
| DIVALIKE+J | -282.5 | 3 | 0.001 | 2.10E-10 | 0.01 | 571.1 | 1.10E-06 |
| **BAYAREALIKE** | **-268.8** | **3** | **0.0005** | **0.0002** | **0.017** | **543.6** | **1** |
| BAYAREALIKE+J | -278 | 3 | 0.0005 | 0.0004 | 0.0072 | 562 | 0.0001 |

**Table S6. Model Fitting for the Historical Biogeographic Reconstruction of the Three Area Dataset.** Bold indicates best-fit model.

| Model | LnL | numparams | d | e | j | AIC | Weight |
|---|---|---|---|---|---|---|---|
| DEC | -146.2 | 2 | 0.0028 | 1.00E-12 | 0 | 296.5 | 1.10E-17 |
| DEC+J | -140.1 | 3 | 0.0023 | 1.00E-12 | 0.017 | 286.3 | 1.80E-15 |
| DIVALIKE | -146.3 | 3 | 0.0027 | 1.00E-12 | 0.013 | 298.6 | 3.80E-18 |
| DIVALIKE+J | -146.3 | 3 | 0.0027 | 1.00E-12 | 0.013 | 298.6 | 3.80E-18 |
| **BAYAREALIKE** | **-106.2** | **3** | **0.0005** | **0.0005** | **0.023** | **218.5** | **0.94** |
| BAYAREALIKE+J | -109 | 3 | 0.0005 | 0.0007 | 0.011 | 224 | 0.06 |

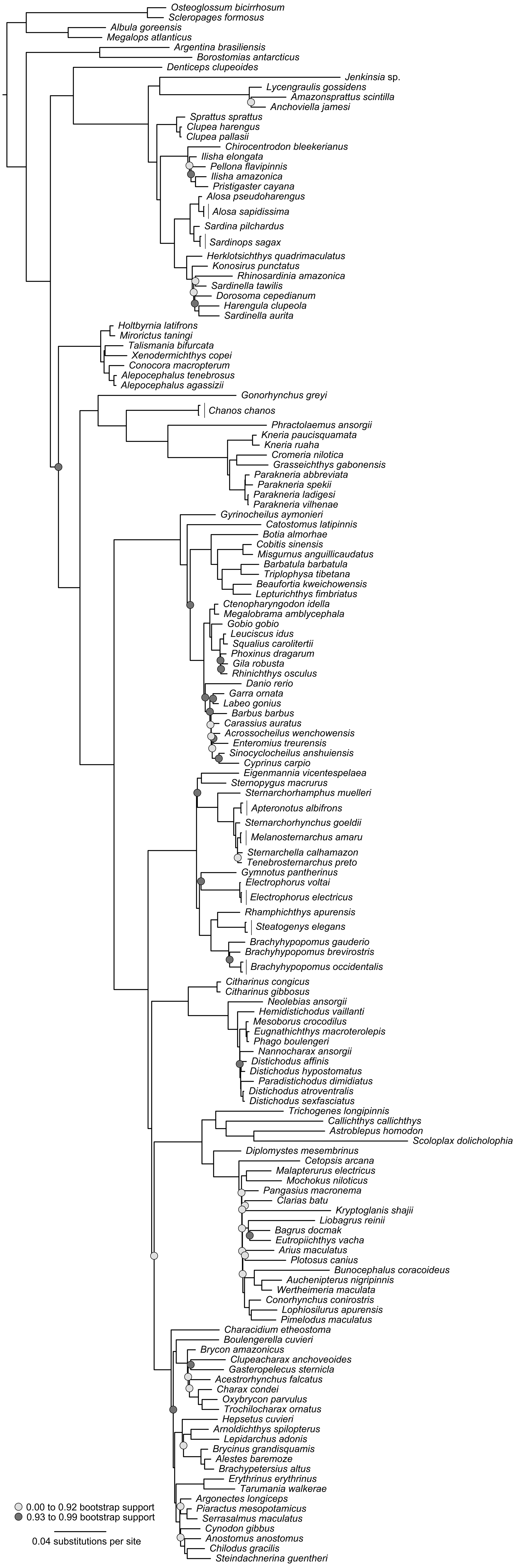
Osteoglossum bicirrhosum
Scleropages formosus
Albula goreensis
Megalops atlanticus
Argentina brasiliensis
Borostomias antarcticus
Denticeps clupeoides
Jenkinsia sp.
Lycengraulis gossidens
Amazonsprattus scintilla
Anchoviella jamesi
Sprattus sprattus
Clupea harengus
Clupea pallasii
Chirocentrodon bleekerianus
Ilisha elongata
Pellona flavipinnis
Ilisha amazonica
Pristigaster cayana
Alosa pseudoharengus
Alosa sapidissima
Sardina pilchardus
Sardinops sagax
Herklotsichthys quadrimaculatus
Konosirus punctatus
Rhinosardinia amazonica
Sardinella tawilis
Dorosoma cepedianum
Harengula clupeola
Sardinella aurita
Holtbyrnia latifrons
Mirorictus taningi
Talismania bifurcata
Xenodermichthys copei
Conocora macropterum
Alepocephalus tenebrosus
Alepocephalus agassizii
Gonorhynchus greyi
Chanos chanos
Phractolaemus ansorgii
Kneria paucisquamata
Kneria ruaha
Cromeria nilotica
Grasseichthys gabonensis
Parakneria abbreviata
Parakneria spekii
Parakneria ladigesi
Parakneria vilhenae
Gyrinocheilus aymonieri
Catostomus latipinnis
Botia almorhae
Cobitis sinensis
Misgurnus anguillicaudatus
Barbatula barbatula
Triplophysa tibetana
Beaufortia kweichowensis
Lepturichthys fimbriatus
Ctenopharyngodon idella
Megalobrama amblycephala
Gobio gobio
Leuciscus idus
Squalius carolitertii
Phoxinus dragarum
Gila robusta
Rhinichthys osculus
Danio rerio
Garra ornata
Labeo gonius
Barbus barbus
Carassius auratus
Acrossocheilus wenchowensis
Enteromius treurensis
Sinocyclocheilus anshuiensis
Cyprinus carpio
Eigenmannia vicentespelaea
Sternopygus macrurus
Sternarchorhamphus muelleri
Apteronotus albifrons
Sternarchorhynchus goeldii
Melanosternarchus amaru
Sternarchella calhamazon
Tenebrosternarchus preto
Gymnotus pantherinus
Electrophorus voltai
Electrophorus electricus
Rhamphichthys apurensis
Steatogenys elegans
Brachyhypopomus gauderio
Brachyhypopomus brevirostris
Brachyhypopomus occidentalis
Citharinus congicus
Citharinus gibbosus
Neolebias ansorgii
Hemidistichodus vaillanti
Mesoborus crocodilus
Eugnathichthys macroterolepis
Phago boulengeri
Nannocharax ansorgii
Distichodus affinis
Distichodus hypostomatus
Paradistichodus dimidiatus
Distichodus atroventralis
Distichodus sexfasciatus
Trichogenes longipinnis
Callichthys callichthys
Astroblepus homodon
Scoloplax dolicholophia
Diplomystes mesembrinus
Cetopsis arcana
Malapterurus electricus
Mochokus niloticus
Pangasius macronema
Clarias batu
Kryptoglanis shajii
Liobagrus reinii
Bagrus docmak
Eutropiichthys vacha
Arius maculatus
Plotosus canius
Bunocephalus coracoideus
Auchenipterus nigripinnis
Wertheimeria maculata
Conorhynchus conirostris
Lophiosilurus apurensis
Pimelodus maculatus
Characidium etheostoma
Boulengerella cuvieri
Brycon amazonicus
Clupeacharax anchoveoides
Gasteropelecus sternicla
Acestrorhynchus falcatus
Charax condei
Oxybrycon parvulus
Trochilocharax ornatus
Hepsetus cuvieri
Arnoldichthys spilopterus
Lepidarchus adonis
Brycinus grandisquamis
Alestes baremoze
Brachypetersius altus
Erythrinus erythrinus
Tarumania walkerae
Argonectes longiceps
Piaractus mesopotamicus
Serrasalmus maculatus
Cynodon gibbus
Anostomus anostomus
Chilodus gracilis
Steindachnerina guentheri
0.00 to 0.92 bootstrap support
0.93 to 0.99 bootstrap support
0.04 substitutions per site

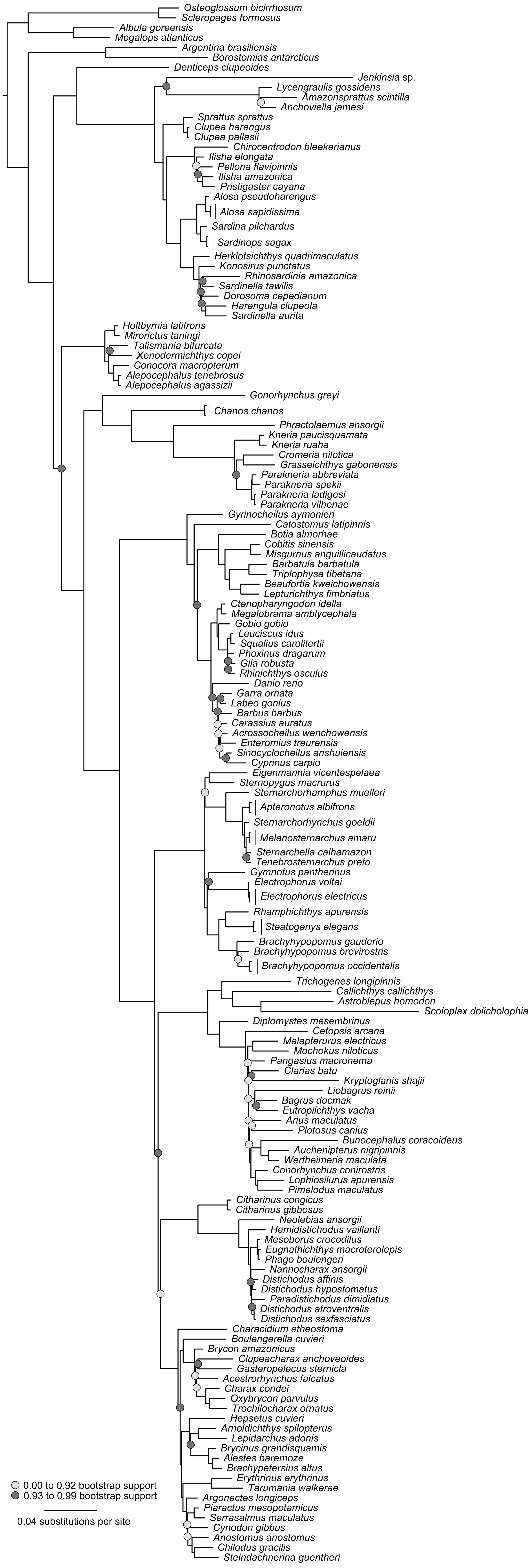

Osteoglossum bicirrhosum
Scleropages formosus
Albula goreensis
Megalops atlanticus
Argentina brasiliensis
Borostomias antarcticus
Denticeps clupeoides
Jenkinsia sp.
Lycengraulis gossidens
Amazonsprattus scintilla
Anchoviella jamesi
Sprattus sprattus
Clupea harengus
Clupea pallasii
Chirocentrodon bleekerianus
Ilisha elongata
Pellona flavipinnis
Ilisha amazonica
Pristigaster cayana
Alosa pseudoharengus
Alosa sapidissima
Sardina pilchardus
Sardinops sagax
Herklotsichthys quadrimaculatus
Konosirus punctatus
Rhinosardinia amazonica
Sardinella tawilis
Dorosoma cepedianum
Harengula clupeola
Sardinella aurita
Holtbyrnia latifrons
Mirorictus taningi
Talismania bifurcata
Xenodermichthys copei
Conocora macropterum
Alepocephalus tenebrosus
Alepocephalus agassizii
Gonorhynchus greyi
Chanos chanos
Phractolaemus ansorgii
Kneria paucisquamata
Kneria ruaha
Cromeria nilotica
Grasseichthys gabonensis
Parakneria abbreviata
Parakneria spekii
Parakneria ladigesi
Parakneria vilhenae
Gyrinocheilus aymonieri
Catostomus latipinnis
Botia almorhae
Cobitis sinensis
Misgurnus anguillicaudatus
Barbatula barbatula
Triplophysa tibetana
Beaufortia kweichowensis
Lepturichthys fimbriatus
Ctenopharyngodon idella
Megalobrama amblycephala
Gobio gobio
Leuciscus idus
Squalius carolitertii
Phoxinus dragarum
Gila robusta
Rhinichthys osculus
Danio rerio
Garra ornata
Labeo gonius
Barbus barbus
Carassius auratus
Acrossocheilus wenchowensis
Enteromius treurensis
Sinocyclocheilus anshuiensis
Cyprinus carpio
Eigenmannia vicentespelaea
Sternopygus macrurus
Sternarchorhamphus muelleri
Apteronotus albifrons
Sternarchorhynchus goeldii
Melanosternarchus amaru
Sternarchella calhamazon
Tenebrosternarchus preto
Gymnotus pantherinus
Electrophorus voltai
Electrophorus electricus
Rhamphichthys apurensis
Steatogenys elegans
Brachyhypopomus gauderio
Brachyhypopomus brevirostris
Brachyhypopomus occidentalis
Trichogenes longipinnis
Callichthys callichthys
Astroblepus homodon
Scoloplax dolicholophia
Diplomystes mesembrinus
Cetopsis arcana
Malapterurus electricus
Mochokus niloticus
Pangasius macronema
Clarias batu
Kryptoglanis shajii
Liobagrus reinii
Bagrus docmak
Eutropiichthys vacha
Arius maculatus
Plotosus canius
Bunocephalus coracoideus
Auchenipterus nigripinnis
Wertheimeria maculata
Conorhynchus conirostris
Lophiosilurus apurensis
Pimelodus maculatus
Citharinus congicus
Citharinus gibbosus
Neolebias ansorgii
Hemidistichodus vaillanti
Mesoborus crocodilus
Eugnathichthys macroterolepis
Phago boulengeri
Nannocharax ansorgii
Distichodus affinis
Distichodus hypostomatus
Paradistichodus dimidiatus
Distichodus atroventralis
Distichodus sexfasciatus
Characidium etheostoma
Boulengerella cuvieri
Brycon amazonicus
Clupeacharax anchoveoides
Gasteropelecus sternicla
Acestrorhynchus falcatus
Charax condei
Oxybrycon parvulus
Trochilocharax ornatus
Hepsetus cuvieri
Arnoldichthys spilopterus
Lepidarchus adonis
Brycinus grandisquamis
Alestes baremoze
Brachypetersius altus
Erythrinus erythrinus
Tarumania walkerae
Argonectes longiceps
Piaractus mesopotamicus
Serrasalmus maculatus
Cynodon gibbus
Anostomus anostomus
Chilodus gracilis
Steindachnerina guentheri
0.00 to 0.92 bootstrap support
0.93 to 0.99 bootstrap support
0.04 substitutions per site

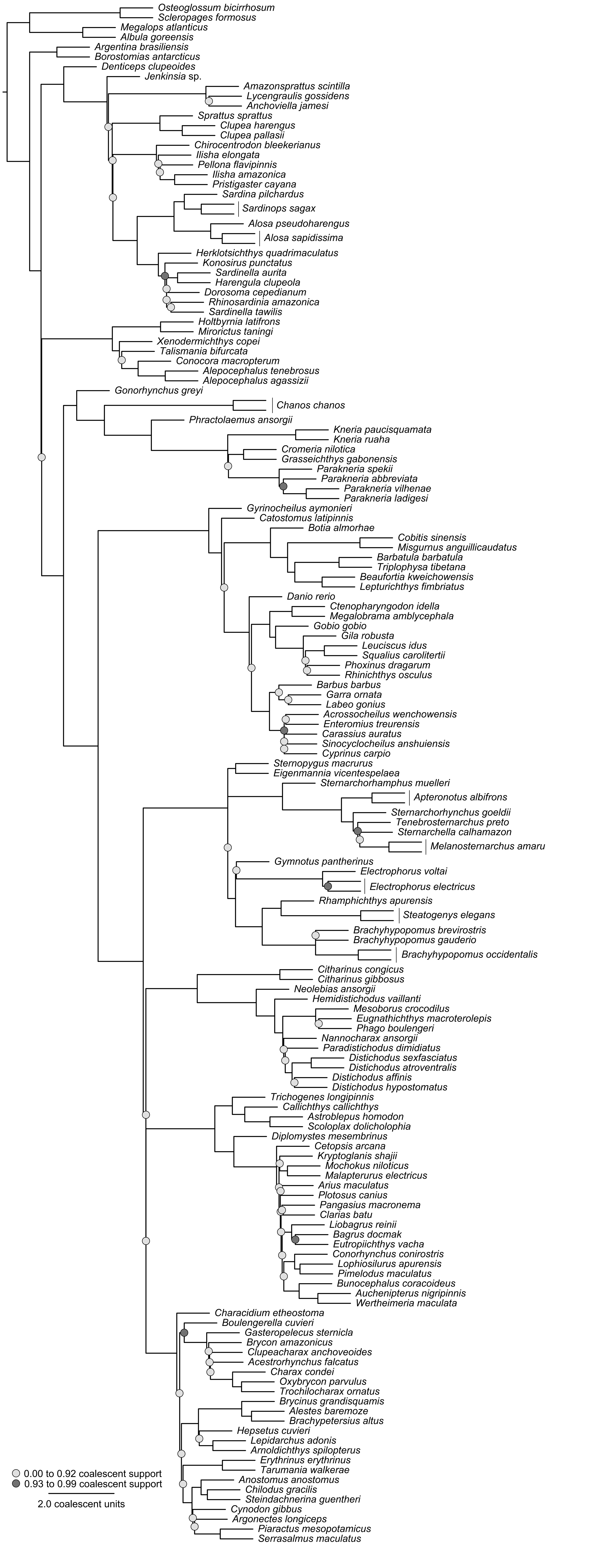
Osteoglossum bicirrhosum
Scleropages formosus
Megalops atlanticus
Albula goreensis
Argentina brasiliensis
Borostomias antarcticus
Denticeps clupeoides
Jenkinsia sp.
Amazonsprattus scintilla
Lycengraulis gossidens
Anchoviella jamesi
Sprattus sprattus
Clupea harengus
Clupea pallasii
Chirocentrodon bleekerianus
Ilisha elongata
Pellona flavipinnis
Ilisha amazonica
Pristigaster cayana
Sardina pilchardus
Sardinops sagax
Alosa pseudoharengus
Alosa sapidissima
Herklotsichthys quadrimaculatus
Konosirus punctatus
Sardinella aurita
Harengula clupeola
Dorosoma cepedianum
Rhinosardinia amazonica
Sardinella tawilis
Holtbyrnia latifrons
Mirorictus taningi
Xenodermichthys copei
Talismania bifurcata
Conocora macropterum
Alepocephalus tenebrosus
Alepocephalus agassizii
Gonorhynchus greyi
Chanos chanos
Phractolaemus ansorgii
Kneria paucisquamata
Kneria ruaha
Cromeria nilotica
Grasseichthys gabonensis
Parakneria spekii
Parakneria abbreviata
Parakneria vilhenae
Parakneria ladigesi
Gyrinocheilus aymonieri
Catostomus latipinnis
Botia almorhae
Cobitis sinensis
Misgurnus anguillicaudatus
Barbatula barbatula
Triplophysa tibetana
Beaufortia kweichowensis
Lepturichthys fimbriatus
Danio rerio
Ctenopharyngodon idella
Megalobrama amblycephala
Gobio gobio
Gila robusta
Leuciscus idus
Squalius carolitertii
Phoxinus dragarum
Rhinichthys osculus
Barbus barbus
Garra ornata
Labeo gonius
Acrossocheilus wenchowensis
Enteromius treurensis
Carassius auratus
Sinocyclocheilus anshuiensis
Cyprinus carpio
Sternopygus macrurus
Eigenmannia vicentespelaea
Sternarchorhamphus muelleri
Apteronotus albifrons
Sternarchorhynchus goeldii
Tenebrosternarchus preto
Sternarchella calhamazon
Melanosternarchus amaru
Gymnotus pantherinus
Electrophorus voltai
Electrophorus electricus
Rhamphichthys apurensis
Steatogenys elegans
Brachyhypopomus brevirostris
Brachyhypopomus gauderio
Brachyhypopomus occidentalis
Citharinus congicus
Citharinus gibbosus
Neolebias ansorgii
Hemidistichodus vaillanti
Mesoborus crocodilus
Eugnathichthys macroterolepis
Phago boulengeri
Nannocharax ansorgii
Paradistichodus dimidiatus
Distichodus sexfasciatus
Distichodus atroventralis
Distichodus affinis
Distichodus hypostomatus
Trichogenes longipinnis
Callichthys callichthys
Astroblepus homodon
Scoloplax dolicholophia
Diplomystes mesembrinus
Cetopsis arcana
Kryptoglanis shajii
Mochokus niloticus
Malapterurus electricus
Arius maculatus
Plotosus canius
Pangasius macronema
Clarias batu
Liobagrus reinii
Bagrus docmak
Eutropiichthys vacha
Conorhynchus conirostris
Lophiosilurus apurensis
Pimelodus maculatus
Bunocephalus coracoideus
Auchenipterus nigripinnis
Wertheimeria maculata
Characidium etheostoma
Boulengerella cuvieri
Gasteropelecus sternicla
Brycon amazonicus
Clupeacharax anchoveoides
Acestrorhynchus falcatus
Charax condei
Oxybrycon parvulus
Trochilocharax ornatus
Brycinus grandisquamis
Alestes baremoze
Brachypetersius altus
Hepsetus cuvieri
Lepidarchus adonis
Arnoldichthys spilopterus
Erythrinus erythrinus
Tarumania walkerae
Anostomus anostomus
Chilodus gracilis
Steindachnerina guentheri
Cynodon gibbus
Argonectes longiceps
Piaractus mesopotamicus
Serrasalmus maculatus
0.00 to 0.92 coalescent support
0.93 to 0.99 coalescent support
2.0 coalescent units

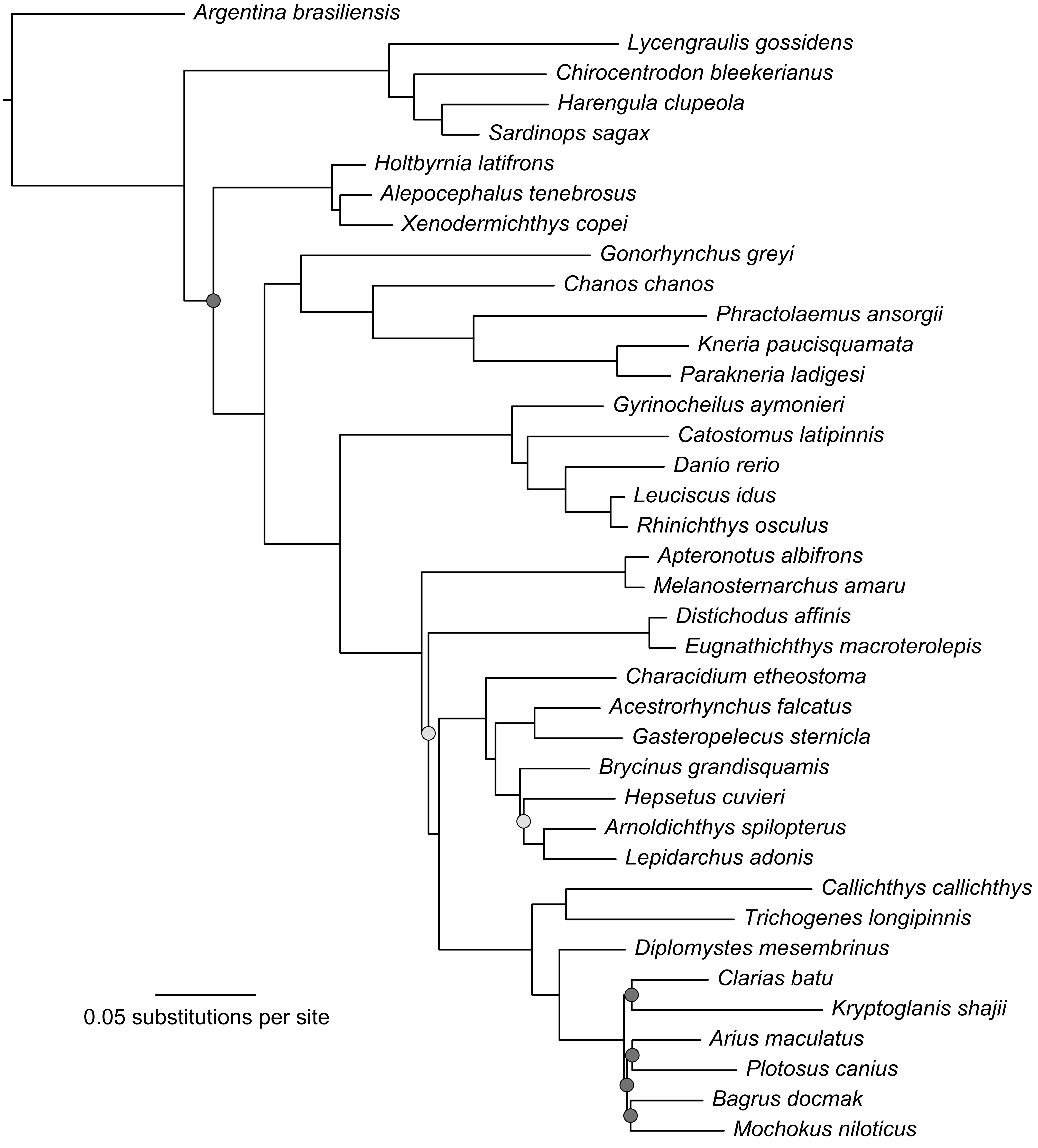

Argentina brasiliensis
Lycengraulis gossidens
Chirocentrodon bleekerianus
Harengula clupeola
Sardinops sagax
Holtbyrnia latifrons
Alepocephalus tenebrosus
Xenodermichthys copei
Gonorhynchus greyi
Chanos chanos
Phractolaemus ansorgii
Kneria paucisquamata
Parakneria ladigesi
Gyrinocheilus aymonieri
Catostomus latipinnis
Danio rerio
Leuciscus idus
Rhinichthys osculus
Apteronotus albifrons
Melanosternarchus amaru
Distichodus affinis
Eugnathichthys macroterolepis
Characidium etheostoma
Acestrorhynchus falcatus
Gasteropelecus sternicla
Brycinus grandisquamis
Hepsetus cuvieri
Arnoldichthys spilopterus
Lepidarchus adonis
Callichthys callichthys
Trichogenes longipinnis
Diplomystes mesembrinus
Clarias batu
Kryptoglanis shajii
Arius maculatus
Plotosus canius
Bagrus docmak
Mochokus niloticus
0.05 substitutions per site

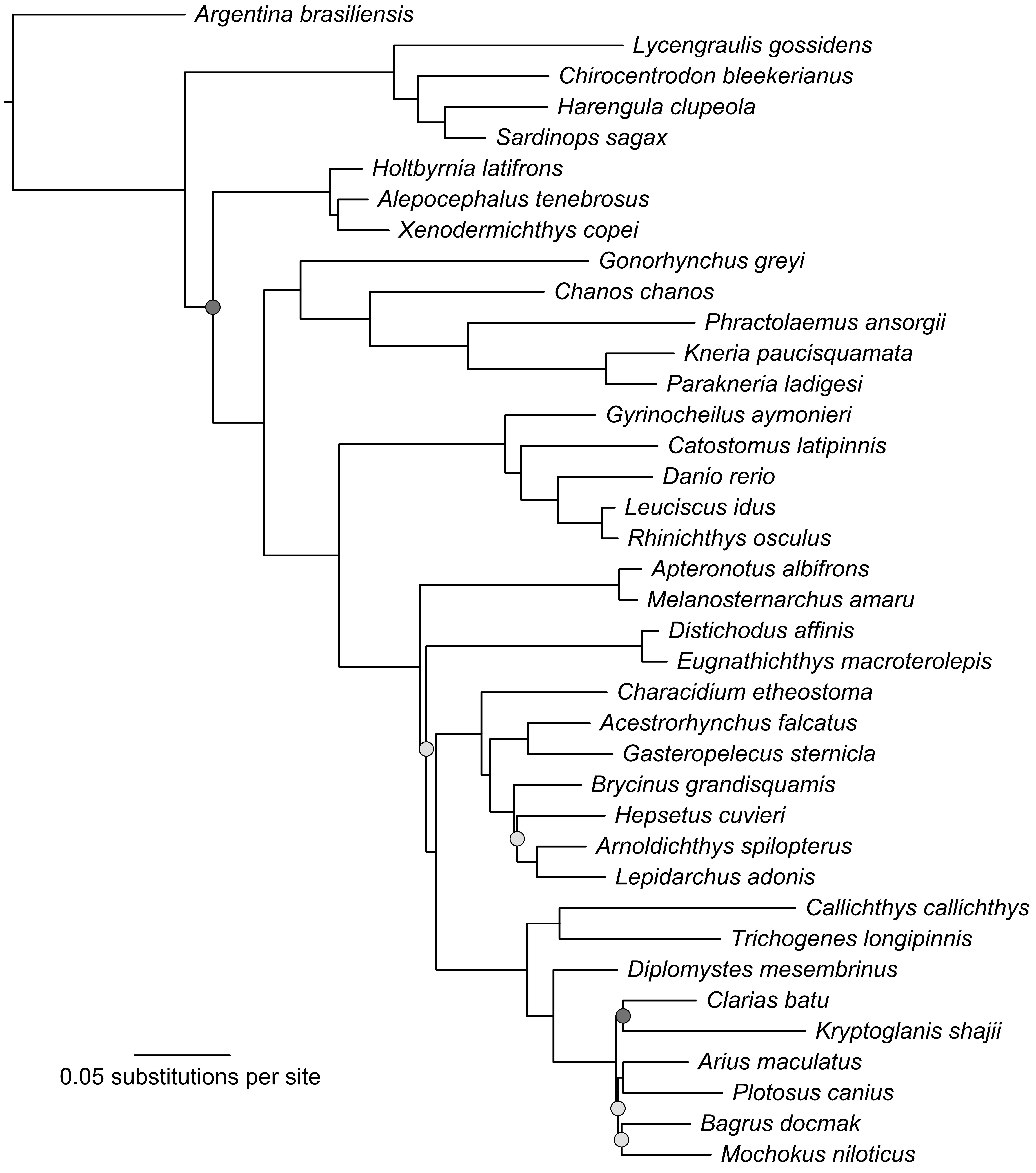

Argentina brasiliensis
Lycengraulis gossidens
Chirocentrodon bleekerianus
Harengula clupeola
Sardinops sagax
Holtbyrnia latifrons
Alepocephalus tenebrosus
Xenodermichthys copei
Gonorhynchus greyi
Chanos chanos
Phractolaemus ansorgii
Kneria paucisquamata
Parakneria ladigesi
Gyrinocheilus aymonieri
Catostomus latipinnis
Danio rerio
Leuciscus idus
Rhinichthys osculus
Apteronotus albifrons
Melanosternarchus amaru
Distichodus affinis
Eugnathichthys macroterolepis
Characidium etheostoma
Acestrorhynchus falcatus
Gasteropelecus sternicla
Brycinus grandisquamis
Hepsetus cuvieri
Arnoldichthys spilopterus
Lepidarchus adonis
Callichthys callichthys
Trichogenes longipinnis
Diplomystes mesembrinus
Clarias batu
Kryptoglanis shajii
Arius maculatus
Plotosus canius
Bagrus docmak
Mochokus niloticus
0.05 substitutions per site

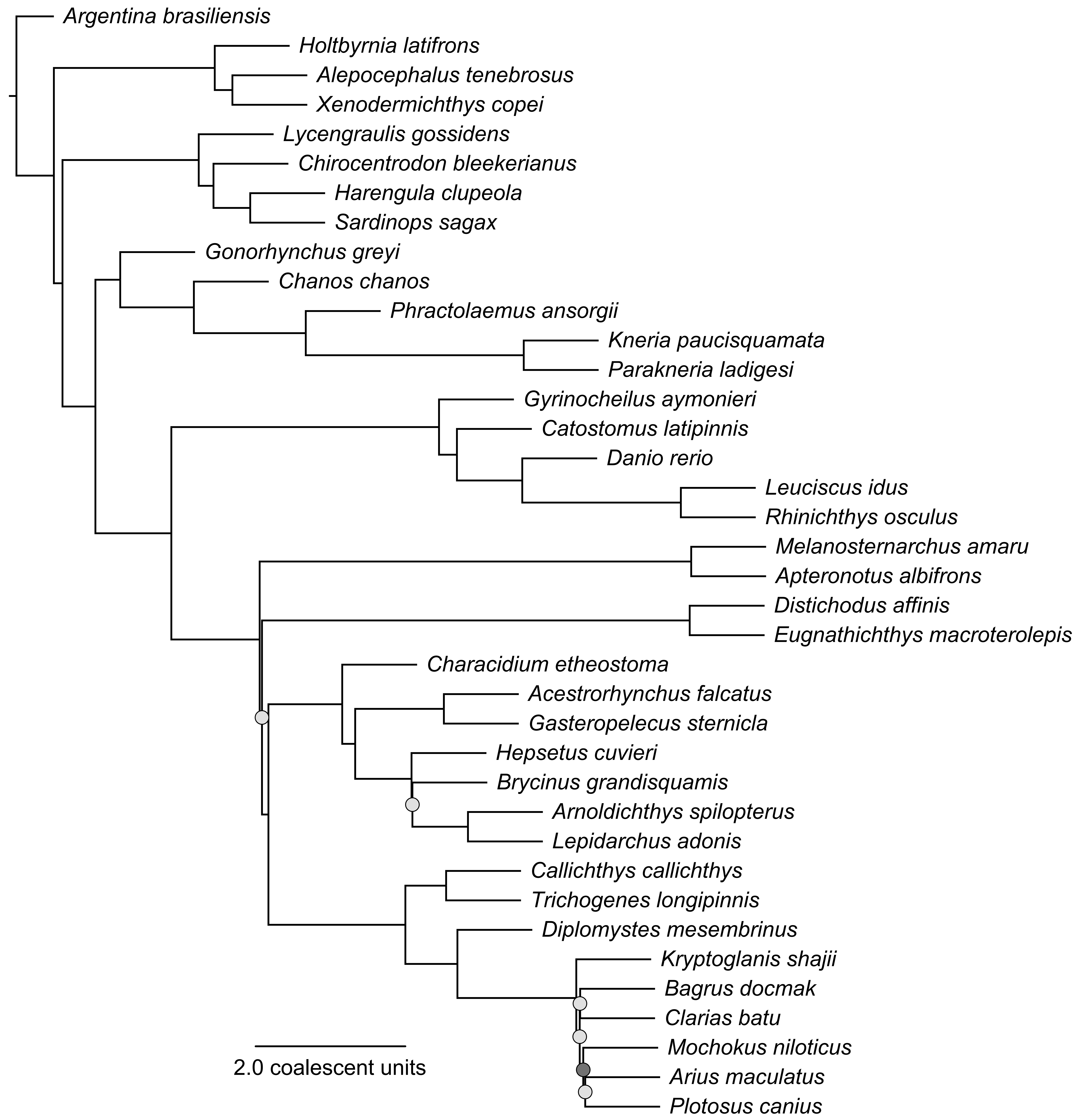
Argentina brasiliensis
Holtbyrnia latifrons
Alepocephalus tenebrosus
Xenodermichthys copei
Lycengraulis gossidens
Chirocentrodon bleekerianus
Harengula clupeola
Sardinops sagax
Gonorhynchus greyi
Chanos chanos
Phractolaemus ansorgii
Kneria paucisquamata
Parakneria ladigesi
Gyrinocheilus aymonieri
Catostomus latipinnis
Danio rerio
Leuciscus idus
Rhinichthys osculus
Melanosternarchus amaru
Apteronotus albifrons
Distichodus affinis
Eugnathichthys macroterolepis
Characidium etheostoma
Acestrorhynchus falcatus
Gasteropelecus sternicla
Hepsetus cuvieri
Brycinus grandisquamis
Arnoldichthys spilopterus
Lepidarchus adonis
Callichthys callichthys
Trichogenes longipinnis
Diplomystes mesembrinus
Kryptoglanis shajii
Bagrus docmak
Clarias batu
Mochokus niloticus
Arius maculatus
Plotosus canius
2.0 coalescent units

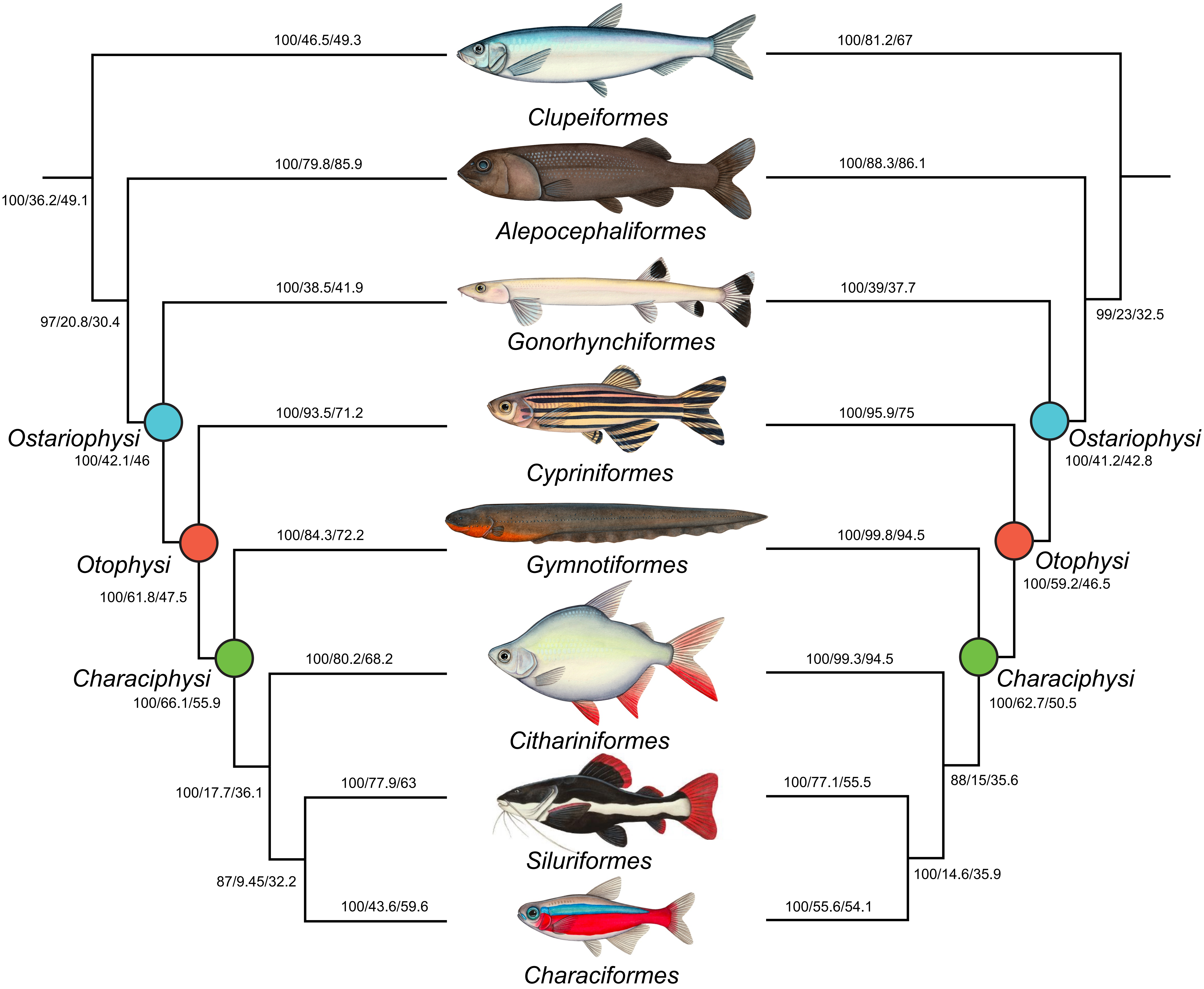
100/46.5/49.3
100/81.2/67
Clupeiformes
100/79.8/85.9
100/88.3/86.1
100/36.2/49.1
Alepocephaliformes
100/38.5/41.9
100/39/37.7
97/20.8/30.4
99/23/32.5
Gonorhynchiformes
100/93.5/71.2
100/95.9/75
Ostariophysi
Ostariophysi
100/42.1/46
100/41.2/42.8
Cypriniformes
100/84.3/72.2
100/99.8/94.5
Otophysi
Gymnotiformes
Otophysi
100/61.8/47.5
100/59.2/46.5
100/80.2/68.2
100/99.3/94.5
Characiphysi
Characiphysi
100/66.1/55.9
100/62.7/50.5
Citharinformes
100/17.7/36.1
100/77.9/63
100/77.1/55.5
88/15/35.6
Siluriformes
87/9.45/32.2
100/14.6/35.9
100/43.6/59.6
100/55.6/54.1
Characiformes

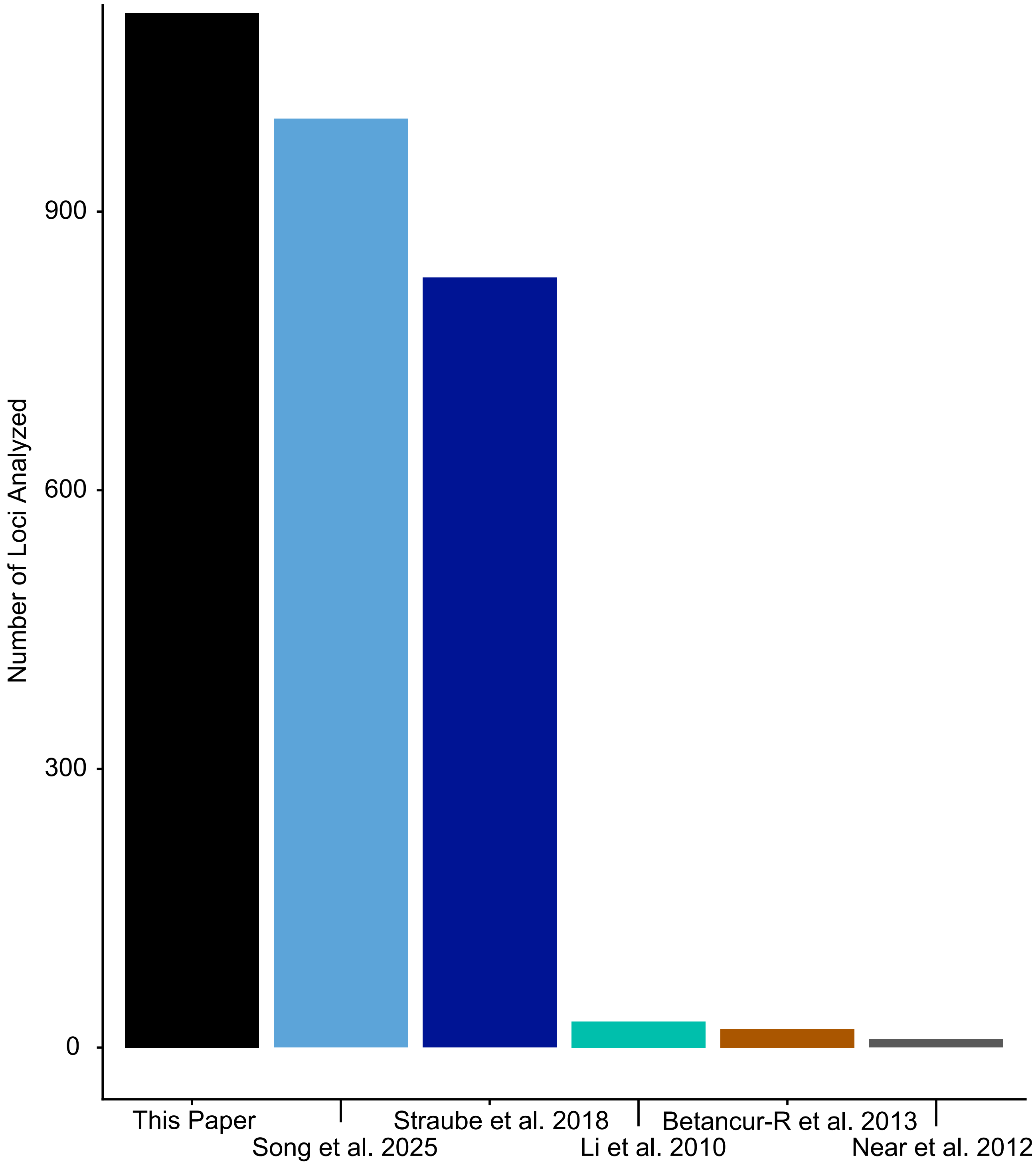

Number of Loci Analyzed
900
600
300
0
This Paper
Song et al. 2025
Straube et al. 2018
Li et al. 2010
Betancur-R et al. 2013
Near et al. 2012

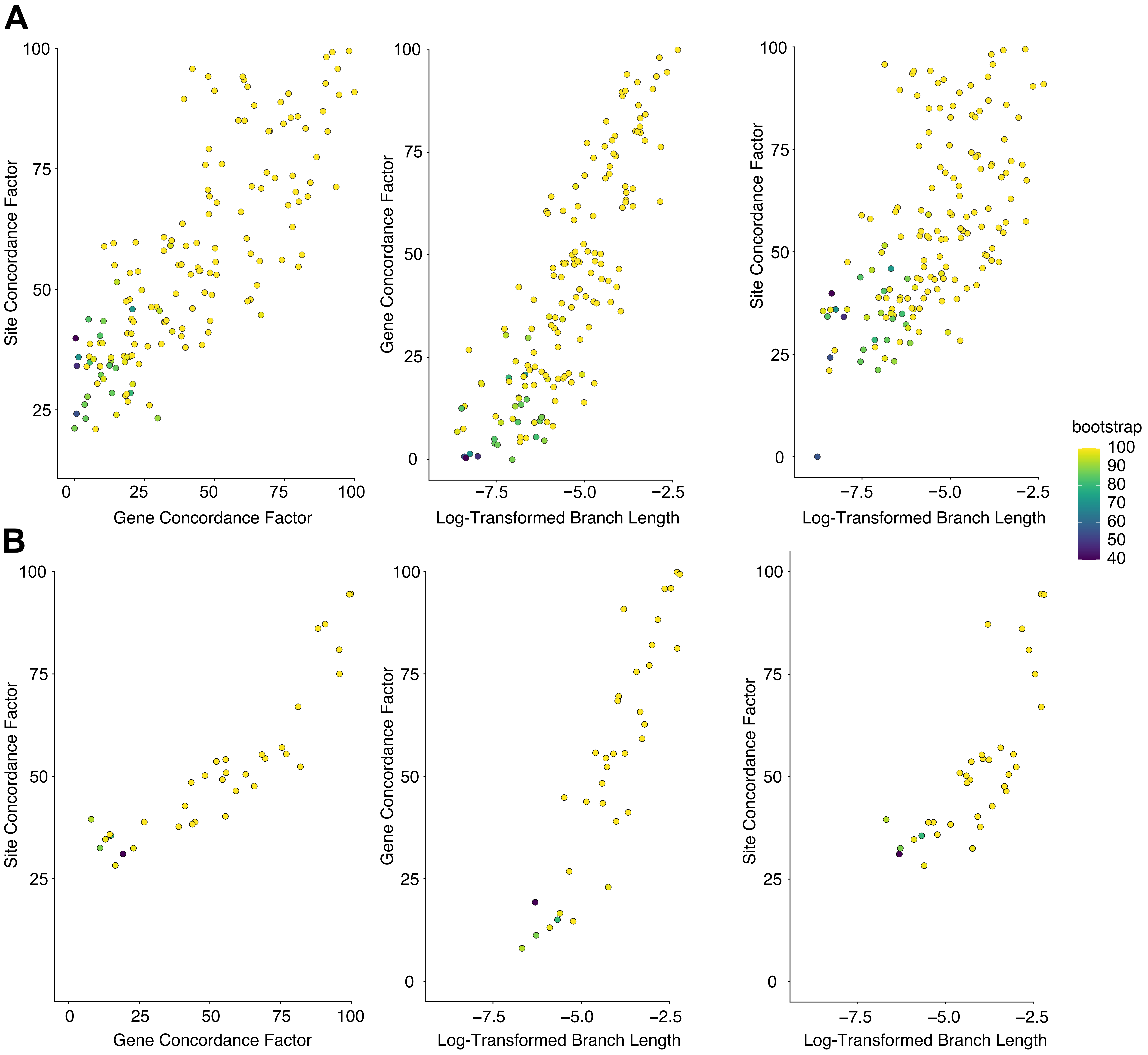
A
B
Site Concordance Factor
Gene Concordance Factor
Log-Transformed Branch Length
bootstrap
100
90
80
70
60
50
40

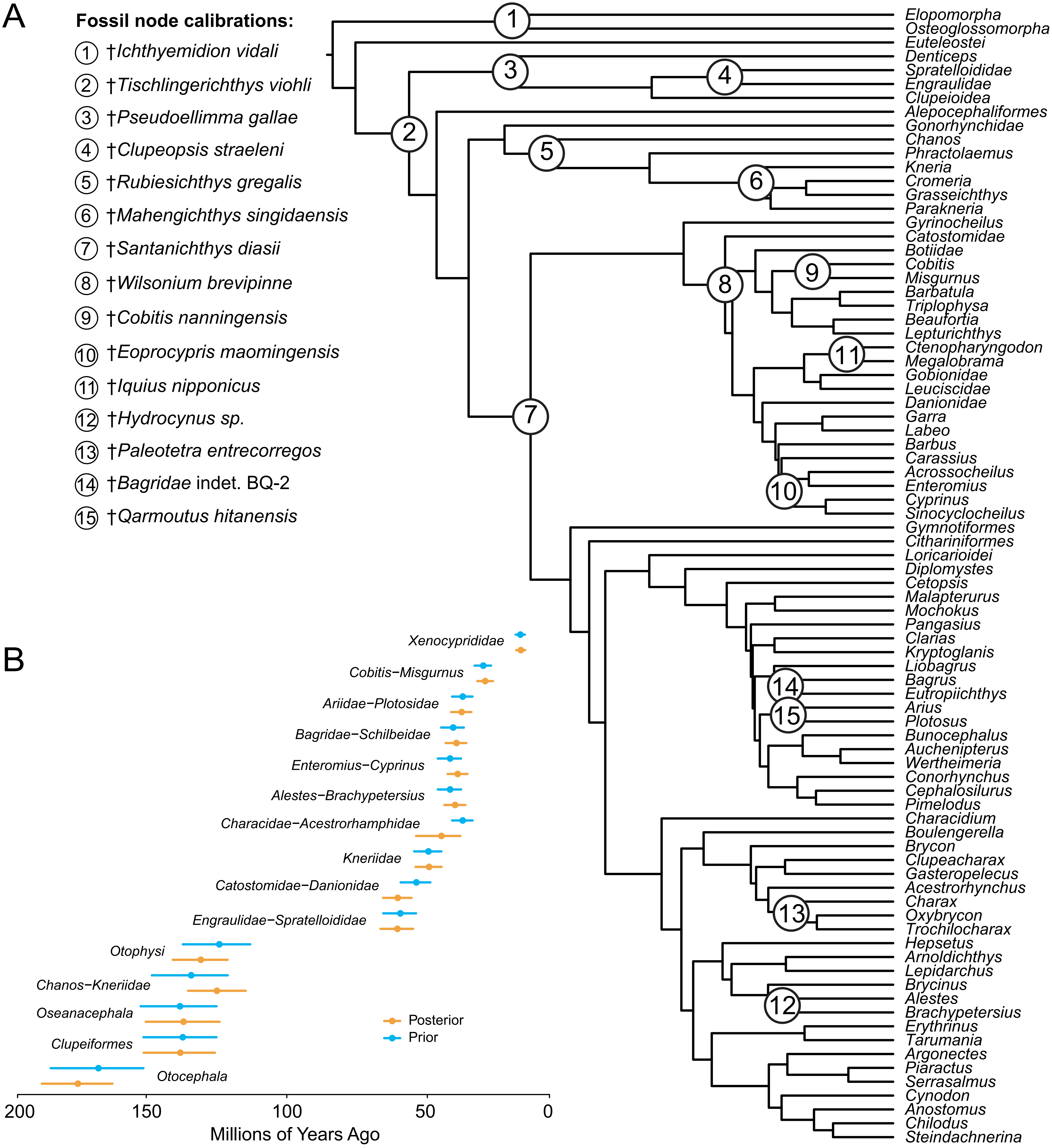
A
Fossil node calibrations:
1 †Ichthyemidion vidali
2 †Tischlingerichthys viohli
3 †Pseudoellimma gallae
4 †Clupeopsis straeleni
5 †Rubiesichthys gregalis
6 †Mahengichthys singidaensis
7 †Santanichthys diasii
8 †Wilsonium brevipinne
9 †Cobitis nanningensis
10 †Eoprocypris maomingensis
11 †Iquius nipponicus
12 †Hydrocynus sp.
13 †Paleotetra entrecorregos
14 †Bagridae indet. BQ-2
15 †Qarmoutus hitanensis
Elopomorpha
Osteoglossomorpha
Euteleostei
Denticeps
Spratelloididae
Engraulidae
Clupeioidea
Alepocephaliformes
Gonorhynchidae
Chanos
Phractolaemus
Kneria
Cromeria
Grasseichthys
Parakneria
Gyrinocheilus
Catostomidae
Botiidae
Cobitis
Misgurnus
Barbatula
Triplophysa
Beaufortia
Lepturichthys
Ctenopharyngodon
Megalobrama
Gobionidae
Leuciscidae
Danionidae
Garra
Labeo
Barbus
Carassius
Acrossocheilus
Enteromius
Cyprinus
Sinocyclocheilus
Gymnotiformes
Citharinioformes
Loricarioidei
Diplomystes
Cetopsis
Malapterurus
Mochokus
Pangasius
Clarias
Kryptoglanis
Liobagrus
Bagrus
Eutropiichthys
Arius
Plotosus
Bunocephalus
Auchenipterus
Wertheimeria
Conorhynchus
Cephalosilurus
Pimelodus
Characidium
Boulengerella
Brycon
Clupeacharax
Gasteropelecus
Acestrorhynchus
Charax
Oxybrycon
Trochilocharax
Hepsetus
Arnoldichthys
Lepidarchus
Brycinus
Alestes
Brachypetersius
Erythrinus
Tarumania
Argonectes
Piaractus
Serrasalmus
Cynodon
Anostomus
Chilodus
Steindachnerina
B
Xenocyprididae
Cobitis−Misgurnus
Ariidae−Plotosidae
Bagridae−Schilbeidae
Enteromius−Cyprinus
Alestes−Brachypetersius
Characidae−Acestrorhamphidae
Kneriidae
Catostomidae−Danionidae
Engraulidae−Spratelloididae
Otophysi
Chanos−Kneriidae
Oseanacephala
Clupeiformes
Otocephala
Posterior
Prior
200
150
100
50
0
Millions of Years Ago

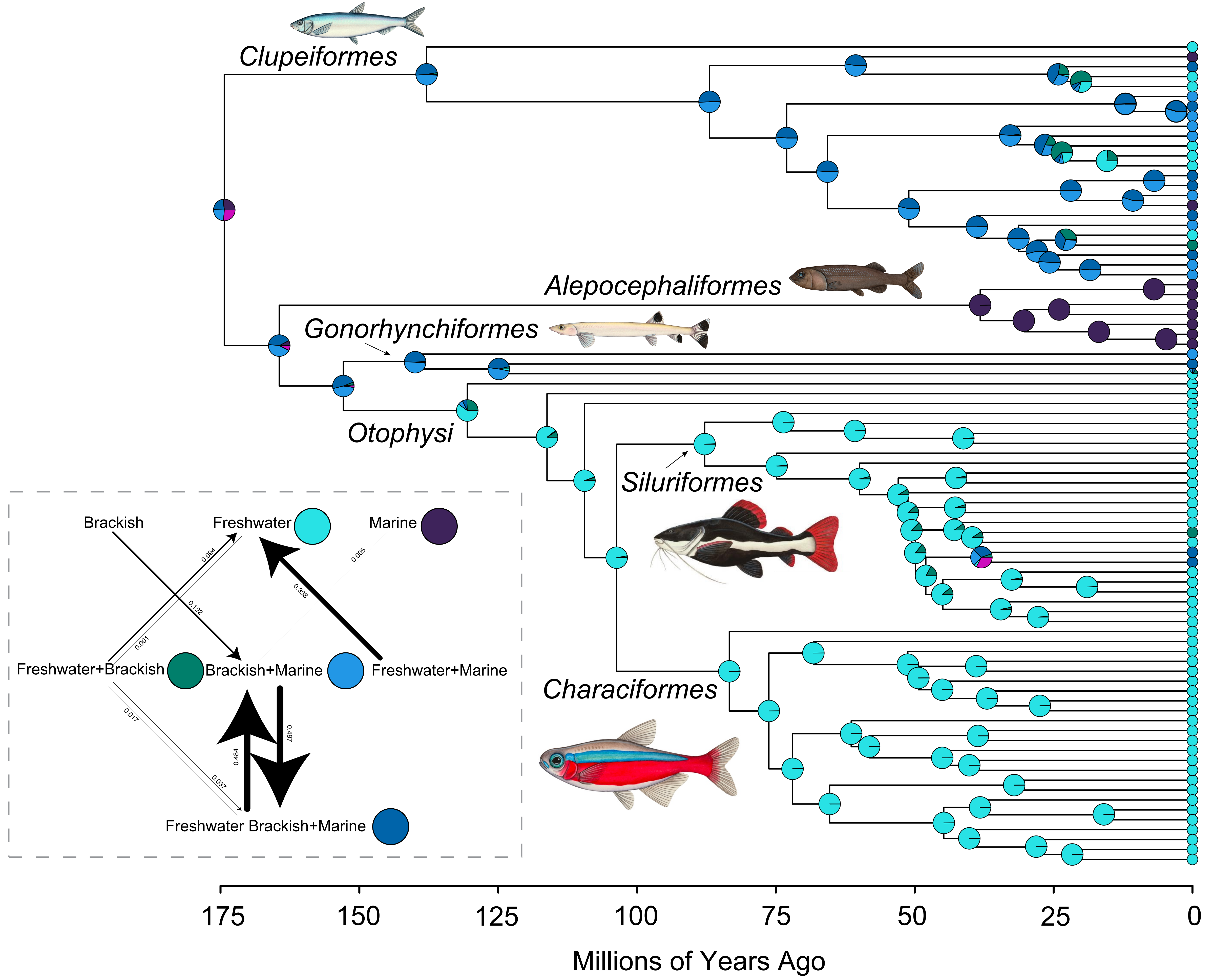
Clupeiformes
Alepocephaliformes
Gonorhynchiformes
Otophysi
Siluriformes
Characiformes
Brackish
Freshwater
Marine
Freshwater+Brackish
Brackish+Marine
Freshwater+Marine
Freshwater Brackish+Marine
0.094
0.122
0.001
0.005
0.338
0.017
0.037
0.484
0.487
175
150
125
100
75
50
25
0
Millions of Years Ago

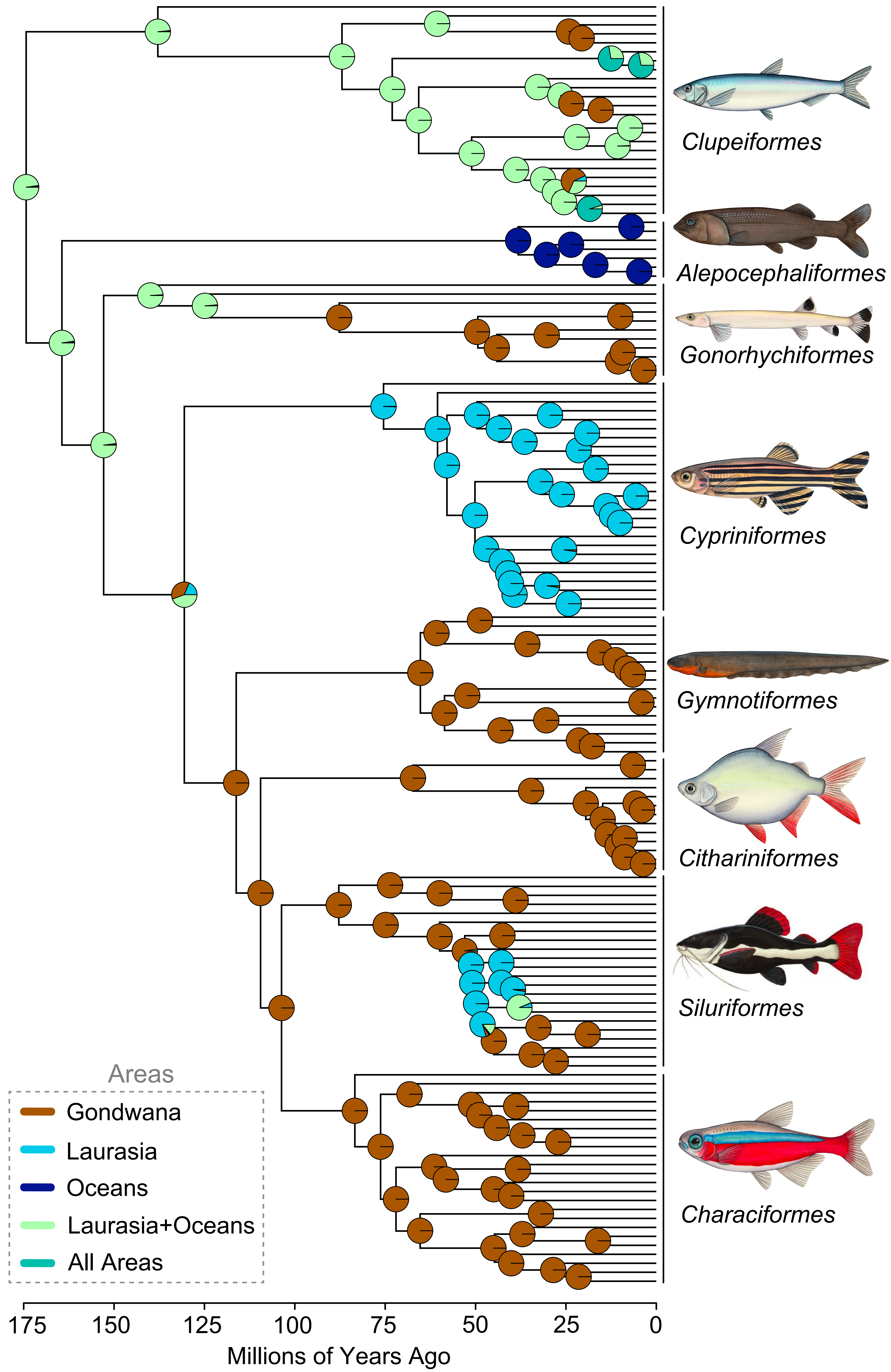

Clupeiformes
Alepocephaliformes
Gonorhychiformes
Cypriniformes
Gymnotiformes
Citharíniformes
Siluriformes
Characiformes
Areas
Gondwana
Laurasia
Oceans
Laurasia+Oceans
All Areas
175
150
125
100
75
50
25
0
Millions of Years Ago
